# Quantifying population resistance to climatic variability: The invasive spotted lanternfly grape pest is buffered against temperature extremes in California

**DOI:** 10.1101/2023.12.21.572841

**Authors:** Stephanie M. Lewkiewicz, Benjamin Seibold, Matthew R. Helmus

## Abstract

Temperature time series data are a composition of average trends and stochastic variability that together shape population dynamics. However, models of temperature-dependent species often overlook variability, focusing solely on growth rate under average conditions. When models omit variability, they can inaccurately predict the dynamics that underlie the establishment of invasive pests sensitive to temperature fluctuations. Here, we conduct a stochastic modeling study of spotted lanternfly (*Lycorma delicatula*), a univoltine grape pest, which has invaded grape growing regions of the eastern U.S. due to human transport leading to frequent establishment of populations in urban and suburban areas. As spotted lanternfly continues to be transported to new grape growing regions and climate change alters variability, it is vital to predict its establishment potential. Although it overwinters as diapausing eggs, experiments suggest that diapause is plastic and not necessary for survival. We developed a deterministic stage-age-structured partial differential equation model of diapausing and non-diapausing populations. We derived a new metric quantifying population resistance to climatic variability defined as the level of stochasticity that leads to negative growth compared to average conditions. We simulated growth rate and resistance to variability across a range of average temperature conditions and stochasticity. We then analyzed how variability and diapause interact with survival, fecundity, and development to affect population dynamics. Finally, we estimated establishment potential across all U.S. cities. Diapausing populations were typically more resistant than non-diapausing populations because diapause enhances overwintering egg survival during winter cold waves, while allowing accelerated development and increased fecundity during summer and fall heat waves. Establishment potential is especially underestimated in important grape growing regions of California if models of diapausing populations omit variability. By quantifying population resistance to climatic variability, we gain a fuller understanding of invasive species establishment in today’s stochastic and changing climate.

## 1. Introduction

Resistance is a key attribute in the description of a physical system, encompassing the ability to remain unchanged despite the presence of a perturbing force (Grimm and Wissel, 1997; Van Meerbeek et al., 2021). In dynamical systems theory, for instance, equilibria are classified according to their resistance properties and the behavior of a system can be inferred from the properties of its core structural features (Meyer, 2016; Strogatz, 2019). In population biology, resistance translates into predictions about population establishment (Smith and Thieme, 2011). A resistant population is one that persists despite external stressors that effect underlying vital processes (Lawson et al., 2015; MacArthur, 1955). The effect of variability on resistance has been probed by investigating stochastic growth rates in deterministic models. Theoretical work has found that on average, stochastic growth rates on average are typically lower than the single deterministic growth rate observed under average conditions (Lawler et al., 2009; Lawson et al., 2015; Vindenes et al., 2014). However, most of this work often oversimplifies the complicated and cumulative impacts of time-dependent temperature trajectories on population dynamics, growth, and establishment potential (Doak et al., 2005). This neglect limits understanding of how species-specific vital rate responses lead to their resistance to climatic variability.

Population responses to temperature variability is of mounting interest due to how temperature dependent mortality, fecundity, development and other vital rates respond to climate change (Stott, 2016). Temperature timeseries data are typically modeled as the sum of a deterministic component, which describes average seasonal patterns, and a stochastic component, which describes deviation from the trend (Ye et al., 2013). In temperate climates, the deterministic component is often a sinusoid over a one-year period defined by the annual mean temperature, annual temperature amplitude, and a phase shift parameter that reflects seasonal lag. Noise is generally characterized by parameters defining autocorrelation and variability magnitude (Ali et al., 2013; Bana et al., 2021; Mudelsee, 2014). Climate change has influenced both the deterministic and stochastic components of temperature. Mean temperatures have increased globally by 1°C on average since the pre-industrial era (Schneider et al., 2022). The magnitude of the deviation of monthly mean temperatures from historical, long-term running averages has increased (Bathiany et al., 2018). Climate change has increased autocorrelation and variability magnitude of land-surface temperatures (Di Cecco and Gouhier, 2018) across multiple timescales (Olonscheck et al., 2021; Zhu et al., 2019) leading to increased frequency of heat and cold waves (Lyon et al., 2019).

Increased temperature variability impacts the population dynamics and establishment potential of invasive species (Root et al., 2003), and invasive species often exhibit wide thermal tolerances and are able to establish under more extreme conditions characteristic of climate change (Gu et al., 2023; Mainka and Howard, 2010). Climatic variability can be more important than average trends in determining invasive insect phenology (Guralnick et al., 2023). Temperature is a primary modulator of insect physiology and behavior (Régnière et al., 2012; Zhang et al., 2013), and climate warming supports insect pest populations by accelerating metabolism, feeding, development, and movement. Higher temperatures can thus lead to greater egg-laying rates, overwintering survival, multivoltinism, and a broader invasive range (Schneider et al., 2022; Skendžić et al., 2021).

Here, we simulate the population dynamics of an insect pest through a partial differential equation (PDE) model, which computes population counts by tracking vital rates as they vary with temperature (Buffoni and Pasquali, 2007; Keyfitz and Keyfitz, 1997; Sharpe and Lotka, 1911). Population dynamical modeling with PDEs has been conducted on many pest species (e.g., Gilioli et al., 2017, 2016; Pasquali et al., 2020, 2019). We implement the model using large ensembles of stochastically generated temperature profiles endowed with both trend and noise components, and compute the reproductive number, *R*_0_, as the multiplicative factor by which the population changes in one year for each temperature profile (e.g., Dee et al., 2020). The PDE is of the spotted lanternfly (SLF, *Lycorma delicatula*), an invasive planthopper establishing across the U.S. (De Bona et al., 2023). SLF is known to feed on over 150 plant species present in North America, including grape (Barringer and Ciafré, 2020; Huron and Helmus, 2022). Invaded vineyards have seen reduced production, increased management costs and vine death (Urban and Leach, 2023). Owing to its risk of disrupting the global wine market (Huron et al., 2022), SLF has been the object of modeling seeking to predict establishment potential based on historical spatiotemporal climatic data (Huron et al., 2022; Jones et al., 2022, 2021; Lewkiewicz et al., 2022; Maino et al., 2021; Wakie et al., 2020). However, no modeling has been performed on its population dynamics, growth rate, and establishment potential accounting for temperature variability. Specifically, we ask: 1) Does variability raise or lower annual growth rates? 2) How do the deterministic vital-rate equations of the PDE interact with the deterministic and stochastic temperature parameters to affect growth rate? 3) How resistant are populations to temperature variability? By answering the final question, we developed a new metric of population establishment that incorporates climatic variability.

## 2. Methods

We first describe the PDE model of the SLF life cycle, including the vital rate response curves calibrated with respect to temperature (Fig. 1). Then we descript the stochastic differential equation that models temperature noise, and the metric of population resistance to temperature variability, Z. Next, we develop and apply a workflow (Fig. 2) based on simulations and a series of plots and videos of SLF population dynamics across temperature parameter sweeps. We identify the mechanisms that underlie changes in vital processes between average and variable temperature conditions to highlight how our analysis can help to understand population dynamical outcomes under changing temperature conditions. Finally, we estimate reproductive number (*R*_0_) and Z values for cities across the continental U.S. and discuss what these values mean for SLF establishment potential.

**Fig. 1:**
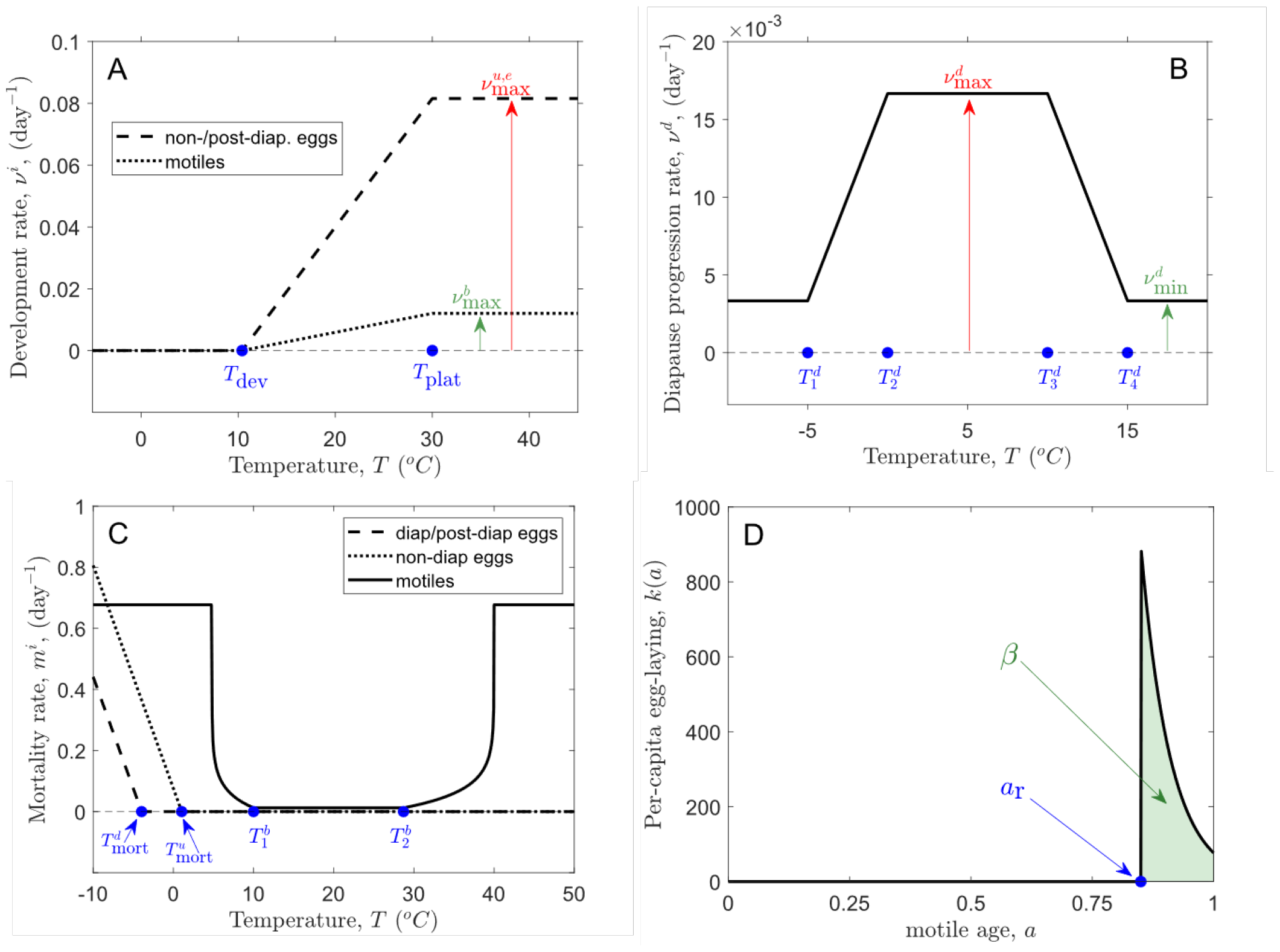
Vital rate functional forms for our spotted lanternfly (Lycorma delicatula) age-structured partial differential equation population dynamic model. The forms are as follows: (A) Degree day function dictating development in eggs and motiles; (B) Rate of egg advection through diapause; (C) Per-capita mortality rate for all life stages; and (D) Density of female eggs laid by female of age a. The area under the curve, β, represents the total number of female eggs laid by one individual over the course of her life. Parameter values are in Table 1. From (A), development rates are zero at temperatures below 10.4°C, increase linearly as temperatures rise from 10.4°C to 30°C, and plateau at the rate v_max_ = 19.6 degree days/day above 30°C. To obtain the development rates specific to each life stage, the degree day function is divided by the length of that life stage in units of degree days (240.3 degree days for developing eggs, 1616.4 degree days for motiles). In (B), diapause advection occurs at its fastest rate of 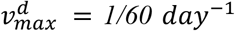 in the optimal temperature range (0°C,10°C), and then decreases from the boundaries of this range, achieving its lowest rate of 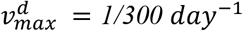 at and below -5°C and above 15°C. In (C), eggs experience cold-induced mortality (basal egg mortality was accounted for in the boundary conditions (Eqs. 2 and 3) through the constant α). Diapause and non-diapause eggs experience cold mortality below -3.96°C, while non-diapause eggs experience mortality below -1.04°C, reflecting the fact that the latter are more vulnerable to cold than the former. The optimal temperature range for motiles is (10°C, 28.7°C), at which they experience a low level of basal mortality, with mortality rates increase at temperatures above and below this range due to heat and cold, respectively. In (D), the egg-laying kernel has an exponentially decaying form across the egg-laying portion of the motile domain, which spans the range (a_r_= 0.85,1).

**Fig. 2:**
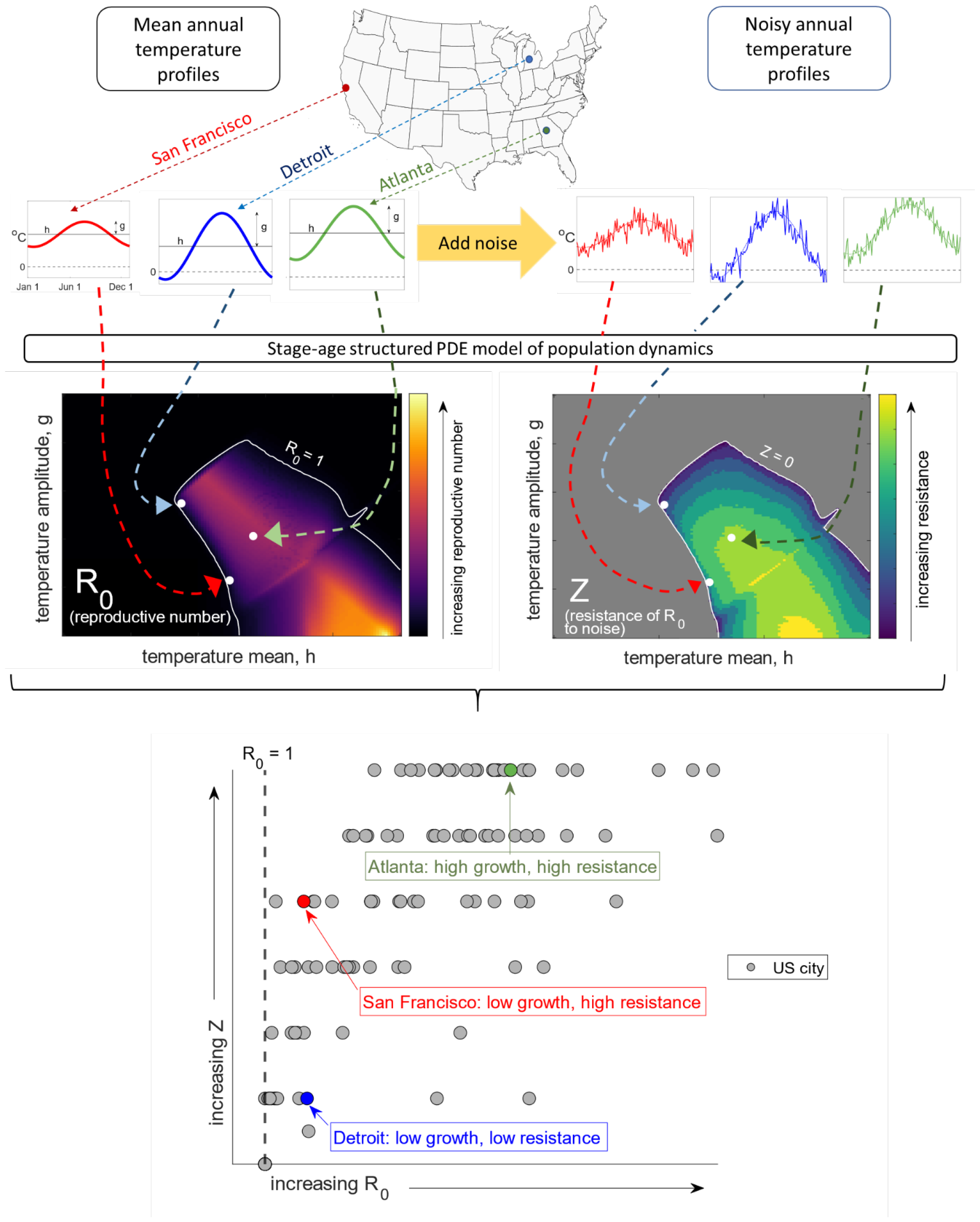
Conceptual workflow of how we estimated the resistance of spotted lanternfly (Lycorma delicatula) population growth and establishment to climate variability. R_0_ is the annual growth factor under mean temperature conditions, represented by sinusoidal temperature profiles with parameters (h, g), the annual mean and amplitude. Historical average temperatures in real United States locations can be fitted to sinusoidal curves to identify where in the (h, g) parameter space the location falls. Population resistance, Z, is the largest noise standard deviation that keeps the median population above equilibrium and is computed using the sinusoidal temperature profiles with added noise. The pairing of an R_0_ and a Z for a given location provides a comprehensive description of establishment potential, including expected behavior and the level of resistance to noise.

### 2.1 Population Dynamics PDE Model

In previous work, we computed the SLF *R*_0_ as a function of the annual mean and amplitude parameters that define the deterministic component of the temperature profile, yielding a comprehensive analysis of growth rates across the full spectrum of possible average temperature profiles (Lewkiewicz et al., 2022). Here, temperature-profile-dependent growth is extended to account for the influence of variability by including the stochastic component of the temperature profile. The average temperature reproductive number is then treated as a baseline measure of expected annual growth, and the nature of the departure from that baseline of growth rates observed under noisy temperature profiles is quantified. We begin with a spatially implicit PDE model of a single SLF population over time. Here, the population is partitioned into four life stages: non-diapause eggs (*ρ*^*u*^), diapausing eggs (*ρ*^*d*^), post-diapause eggs (*ρ*^*e*^), and motiles (*ρ*^*b*^). Diapausing eggs are those currently in diapause, while post-diapause (resp., non-diapause) eggs are those that are developing after having completed (resp., having entirely bypassed) the diapause stage. The motile stage includes juveniles, immature adults, and egg-laying adults. The life stage division was chosen to group individuals with similar parameters of temperature sensitivity (in terms of development, diapause induction/termination, and mortality), regardless of other anatomical and physiological differences between juveniles and adults, or similarities among eggs of different diapause statuses (Lewkiewicz et al., 2022).

**Table 1:** Parameter values of spotted lanternfly PDE model vital rate functions.

| Parameter | Value | Eq. or Fig. | Meaning | Source |
| --- | --- | --- | --- | --- |
| $\sigma^u, \sigma^e$ | $6.4 \times 10^{-4}$ | (1) | diffusion coefficient scaling factor* (developing eggs) | D. Calvin |
| $\sigma^d$ | 0 | (1) | diffusion coefficient scaling factor* (diapause eggs) | --- |
| $\sigma^b$ | $5 \times 10^{-3}$ | (1) | diffusion coefficient scaling factor* (motiles) | D. Calvin |
| $\alpha$ | 0.6 | (2)(3) | egg survival probability | (Keena and Nielsen, 2021; Park, 2015) |
| $T_{dev}$ | 10.4°C | Fig. 1a | development threshold | (Smyers et al., 2021) |
| $v_{max}$ | 19.6 degree days/day | Fig. 1a | maximum development rate | (Smyers et al., 2021) |
| $T_{plat}$ | 30°C | Fig. 1a | maximum development rate threshold | (Smyers et al., 2021) |
| $(T_2^d, T_3^d)$ | (0°C, 10°C) | Fig. 1b | temperature range of maximum diapause advection rate | (Shim and Lee, 2015) |
| $(T_1^d, T_4^d)$ | (−5°C, 15°C) | Fig. 1b | temperature range of elevated diapause advection rate | (Shim and Lee, 2015) |
| $v_{max}^d$ | (1/60) per day | Fig. 1b | maximum diapause advection rate | (Shim and Lee, 2015) |
| $v_{min}^d$ | (1/300) per day | Fig. 1b | minimum diapause advection rate | (Shim and Lee, 2015) |
| $T_{mort}^u$ | 1.04°C | Fig. 1c | cold death temperature threshold (non-diapause eggs) | (Keena and Nielsen, 2021; Park, 2015; Thomas et al., 2012) |
| $T_{mort}^d$ | −3.96°C | Fig. 1c | cold death temperature threshold (diapause lineage eggs) | (Keena and Nielsen, 2021; Park, 2015; Thomas et al., 2012) |
| $(T_1^b, T_2^b)$ | (10°C, 28.7°C) | Fig. 1c | temperature range of basal motile mortality rate | (Kreitman et al., 2021) |
| $a_r$ | 0.85 | Fig. 1d | developmental age of initial egg-laying | D. Calvin |
| $\beta$ | 50 | Fig. 1d | total per-capita<br>lifetime egg-laying | T. Leskey |

Names listed are personal communications from researchers working on spotted lanternfly vital rates.
\*diffusion coefficient scaling factors are necessarily unitless

In the model, diapause is induced in the eggs of fecund females based on the current photoperiod at the time of egg-laying: eggs laid after the summer solstice and before the winter solstice in a given year are assumed to begin diapause when laid, while eggs laid outside of this window are assumed to bypass diapause and begin development immediately when laid. While it is unknown what exactly induces diapause, there is evidence that eggs do not need to go through diapause to hatch and develop (Keena and Nielsen, 2021; Mousseau and Dingle, 1991; Saunders et al., 2004). Eggs progress through and exit diapause at a rate dictated by ambient temperatures, and, upon completion of diapause, undergo development. Post-diapause and non-diapause eggs experience the same temperature-driven development process, but post-diapause eggs have a higher level of cold-tolerance as a consequence of diapause (Thomas et al., 2012). After hatching, juveniles from the diapause and non-diapause lineages enter the same motile pool and are regarded as indistinguishable. Surviving individuals senesce out of the system once they pass through four instar stages, an immature adult stage, and an egg-laying stage (Dara et al., 2015; Liu, 2019).

The dynamics of this system are described by the following advection-diffusion-reaction equations with parameters defined as in Fig. 1 and Table 1:

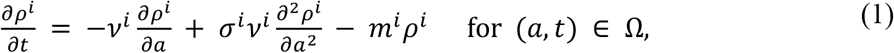

coupled with the following boundary conditions,

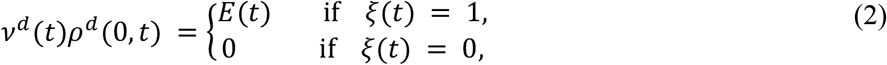

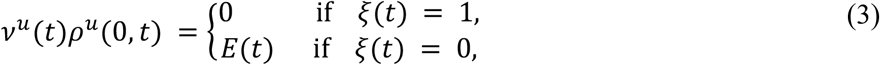

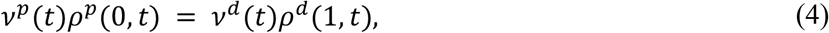

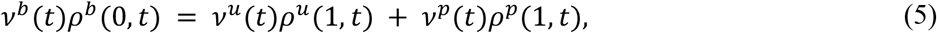

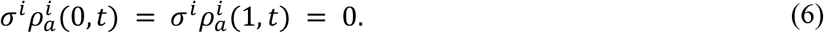

Here, *ρ*^*i*^(*a, t*) is the abundance of individuals of age *a* in life stage *i* at time *t*. The labels *i* = *u, d, e*, and *b* represent non-diapause eggs, diapausing eggs, post-diapause eggs, and motiles, respectively. The variable *a* denotes relative age within a life stage, with ages *a* = 0 and *a* = 1 representing the start and end of the stage. The domain Ω in Eq. (1) is the set {(*a, t*) : *a* ∈ (0, 1), *t* ∈ (0, 365)}, where *t* is measured in days. (We study dynamical behavior on the time scale of one Julian year assumed to have 365 days (Lewkiewicz et al., 2022).) In Eqs. (1-6), *v*^*i*^ denotes the average development rate and *m*^*i*^ the per-capita mortality rate associated with life stage *i*, while *σ*^*i*^ is a diffusion coefficient that accounts for a small amount of interindividual variability in rates of development. Thus, the first two terms on the right-hand side (RHS) of Eq. (1) represent development; the first accounts for average development and the second accounts for the distribution around that average. (Two individuals may develop at slightly different rates due to variation in their individual circumstances, such as exposure to different host species.) The second term on the RHS is scaled by *v*^*i*^ under the assumption that intraspecific variation in development is proportional to the rate of development itself. The third term on the RHS of Eq. (1) is the total mortality rate. The terms *v*^*i*^(*t*)*ρ*^*i*^(0, *t*) and *v*^*i*^ (*t*)*ρ*^*i*^(1, *t*) in Eqs. (2-6) respectively represent the influxes and outfluxes of individuals in life stage *i*. Eqs. (2-6) thus describe life stage transitions by balancing fluxes from one life stage to the next. In Eqs. (2) and (3), 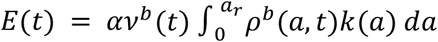 is the rate of egg-laying; *α* is the probability of an egg surviving to hatching under optimal conditions, and *k*(*a*) is the number of eggs laid by a female of age *a*. The function *ξ*(*t*) equals 1 when *t* falls after the summer solstice and before the winter solstice in a given year, and equals 0 otherwise, so that Eqs. (2) and (3) direct eggs into the diapause or non-diapause domains based on seasonality.

The terms *v*^*i*^ and *m*^*i*^ both depend on temperature, *T*, so that *v*^*i*^ *= v*^*i*^(*T*), *m*^*i*^ = *m*^*i*^(*T*), while *σ*^*i*^ is a constant. We take *σ*^*i*^ > 0 for *i* = *u, e*, and *b*, with *σ*^*d*^ = 0, due to a lack of knowledge of interindividual variation in the protein-clearing processes that dictate diapause progression. The exact values of *σ*^*i*^ used in simulations can be found in Table 1, along with all of the other relevant parameter values described in this section.

The developmental rate functions *v*^*i*^(*T*) for *i* = *u, e*, and *b* are rescaled versions of the piecewise-linear SLF degree day rate function introduced in (Smyers et al., 2021). Here, the term “degree day” refers to the *growing* degree day, a unit measuring heat accumulation and, by extension, temperature-dependent development in poikilothermic species (in our case, insect pests). This degree day rate function, which we denote here by *DD*(*T*), assigns to each temperature a certain number of degree days by which an individual advances in developmental age in one day of exposure to that temperature. It therefore has units of “degree days per day,” and the total number of degree days accumulated between two times, *t*_1_ and *t*_2_, is 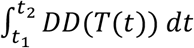. For SLF, no development occurs below *T*_dev_ = 10.4^*o*^C (the development threshold), from which point the development rate increases linearly with *T* until a plateau value of *v*_max_ = 19.6 degree days/day is reached at *T*_plat_ = 30^*o*^C (Smyers et al., 2021). To form the PDE functions *v*^*i*^(*T*) for *i* = *u, e*, and *b, DD*(*T*) is divided by the length of each life stage in degree days to match the unitless age variable, *a*. For developing egg stages (non-diapause and post-diapause eggs), this is 240.3 degree days, while for motiles, it is 1628.4 degree days; in other words, *v*^*i*^ (*T*) = *DD*(*T*)/240.3 for *i* = *u, e*, and *v*^*b*^ (*T*) = *DD*(*T*)/1628.4 (Lewkiewicz et al., 2022). These rescaled functions, depicted in Fig. 1a, thus have units “day^−1^.”

Throughout this paper, we work with and discuss the function *DD*(*T*) when focusing on species biological aspects of our analysis, and with the functions *v*^*i*^(*T*) when focusing on mathematical aspects. For *i* = *d, v*^*d*^(*T*) represents the rate at which eggs progress through diapause. Eggs are expected to progress through diapause at their fastest rate of 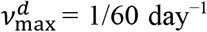 in the optimal cool temperature range 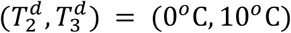 (Fig. 1b) (Shim and Lee, 2015). Eggs move through diapause at their slowest rate, 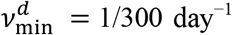, at sufficiently cold and hot temperatures (below 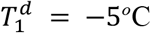 and above 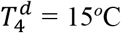) (Shim and Lee, 2015). The mortality functions *m*^*i*^(*T*) for *i* = *u, d*, and *e* are all set to zero above a certain threshold temperature, increasing linearly as temperatures decrease below the mortality threshold. (See Fig. 1c) For *i* = *u*, this mortality threshold is 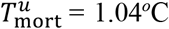, while for *i* = *d* and *e*, the mortality threshold is 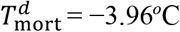, reflecting the assumption that diapause and post-diapause eggs have the same heightened tolerance to cold, which is higher than that experienced by non-diapause eggs (Keena and Nielsen, 2021; Park, 2015). Finally, the function *m*^*b*^(*T*), also depicted in Fig. 1c, features a basal, temperature-independent mortality rate for motiles at mild temperatures 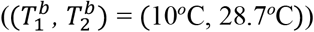, with temperature-induced mortality commencing below 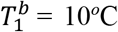 and above 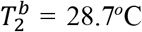 (Kreitman et al., 2021). From the kernel, *k*(*a*), in Fig. 1d, egg-laying begins at motile age *a* = 0.85 (approximately 1616.4 degree days since hatching) and decays exponentially with *a* (Fox, 1993). Further details about the model formulation and calibration to existing SLF data can be found in (Lewkiewicz et al., 2022).

### 2.2 Temperature Profile Model

Seasonal temperature patterns are incorporated into the model (Eqs. (1-6)) with the inclusion of a temperature profile, *T*(*t*), representing temperature over time. In this paper, *T*(*t*) contains a deterministic and a stochastic component; the former accounts for long-term annual and seasonal trends (mean behavior), while the latter incorporates variability and deviation from that average behavior. *T*(*t*) is a stochastic process, with *T*(*t*) = *D*(*t*) + *S*(*t*), where the deterministic component is:

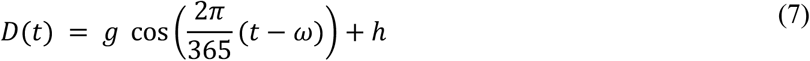

and the stochastic component is:

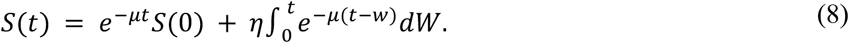

We note that *D*(*t*) is a function of *t*, while *S*(*t*) is a stochastic process. In Eq. (7), *h* is the annual mean temperature, *g*, is the annual temperature amplitude, and *ω* is the phase shift. The phase shift parameter represents the Julian day number of the hottest day of the year. This day varies little across the continental United States, so we used *ω* = 203 (July 22nd) for all of our analyses. The stochastic process in Eq. (8) is the solution of the Langevin equation,

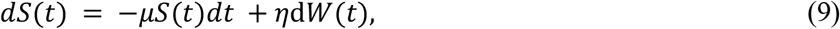

where *μ* and *η* are referred to as the drift and volatility parameters, respectively, and *W*(*t*) is Brownian motion (Evans, 2013). Per Eq. (8), the parameter *μ* dictates the exponentially decaying influence of 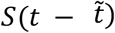 on *S*(*t*), where 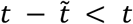, while *η* controls the magnitude of the noise term. From Eq. (8), the influence of 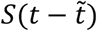 on *S*(*t*) decays exponentially with increasing 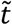. *S*(*t*) has expectation *E*(*S*(*t*)) = *e*^−μ*t*^*E*(*S*(0)) and variance

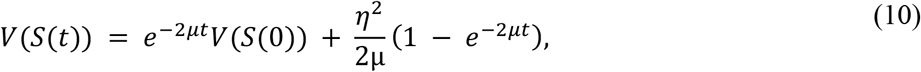

from which it follows that *E*(*S*(*t*)) → 0 and 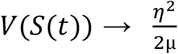 as *t* → ∞ (Evans, 2013). For reasonable choices of *S*(0), the convergences of *E*(*S*(*t*)) and *V*(*S*(*t*)) tend to be rapid, and we may regard *S*(*t*) as having the nearly constant mean zero and variance 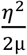 (or standard deviation 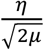). We then define 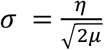 and refer to *σ* as the standard deviation parameter of temperature variability.

In our subsequent analysis, we regarded *T*(*t*) as being defined jointly by the deterministic mean and amplitude parameters, *h* and *g*, and the stochastic parameter, *σ*, with respect to the three of which simulations and parameter sweeps are conducted below. (Numerical tests indicated that simulation outcomes were generally insensitive to the parameter *μ*—which dictated temporal autocorrelation in the noise term—except when |*μ*| ≪ 1, representing unrealistically high levels of autocorrelation. For simplicity, in the rest of the manuscript, we took *μ* = – ln 0.5, under which assumption the influence of *S* at earlier times on *S* at later times decays at a rate of 50% per day.) The parameter sweeps consisted of ensemble simulations of the PDE for large sets of realizations of *T*(*t*); an individual realization of *T*(*t*) was obtained by simply summing the function *D*(*t*) (Eq. (7)) and a realization of the stochastic process *S*(*t*) (Eq. (8)) generated with the Ito integral approximation to Eq. (8) (Evans, 2013).

### 2.3 Population Resistance Metric

In previous work, we computed the SLF reproductive number, *R*_0_, under the purely deterministic temperature model *D*(*t*) of Eq. (7). *D*(*t*) represents an approximation to long-term mean seasonal temperature patterns in a given location, and *R*_0_ is thus the annual growth factor—the factor by which a population grows or declines over the course of one year—under average temperature conditions. Viewing the incorporation of the stochastic process *S*(*t*) (Eq. (8)) into the temperature profile as a disturbance to the system that introduces natural variability (e.g., random fluctuations, cold snaps and heat waves), we developed a metric of how annual growth factors depart from the baseline *R*_0_ as variability intensifies. Specifically, the metric asks: if *R*_0_ > 1 for some (*h, g*) pairing, how large does *σ* need to be to drive growth factors below 1?

To quantify this value of *σ* at which resistance is lost (*R*_0_ < 1) for a fixed set of parameters (*h, g*), we deployed an ensemble simulation in the parameter *σ*. For each value of *σ* in the set {0.1, 0.5, 1, 2, …, 8}, we generated 300 realizations of *S*(*t*) with stochastic parameter *σ*, each of which was added to the same *D*(*t*) (with deterministic parameters (*h, g*)) to produce a set of 300 realizations of *T*(*t*) defined by the three parameters (*h, g, σ*). Precisely, we denote by *S*_*σ,k*_(*t*) the *k*-th realization of *S*(*t*) defined by *σ*, and set *T*_*σ,k*_(*t*) = *D*(*t*) + *S*_*σ,k*_(*t*). Inserting each such *T*_*σ,k*_(*t*) into Eqs. (1-6), the PDE is solved for one year starting on the hottest day of the year using the eigenvector of the one-year numerical solution operator’s dominant eigenvalue as the initial condition. The dominant eigenvector is a natural choice of initial condition in this context because, without the addition of temperature variability, the life stage distribution after one year would be identical in shape to the starting distribution, having only grown or decayed by a factor of *R*_0_. This allows us to observe how growth factors occurring in the presence of temperature variability depart from the baseline growth factor, *R*_0_.

Now denote by 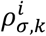 the solution of Eq. (1) using the temperature profile *T*_*σ,k*_(*t*), and define the growth factor, *G*_*σ,k*_, of this individual simulation as,

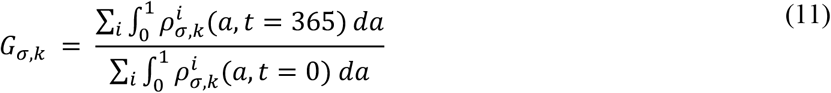

where the summation with respect to *i* is over all four life stages. Eq. (11) is the ratio of the total number of individuals in the population at the end (*t* = 365) versus the start (*t* = 0) of the simulation, thereby representing the factor by which the population has changed in one year.

We define population resistance, *Z*, as,

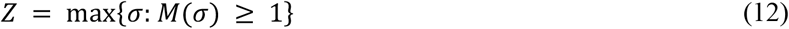

where *M*(*σ*) is the median of the growth factors 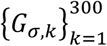.. (The median is considered over the mean because we are interested in the spectrum of possible outcomes of one-year simulations, as opposed to average behavior over many noisy simulations.) The metric *Z* is the largest value of *σ* for which *M*(*σ*) remains at or above 1 i.e., the largest noise standard deviation that keeps the median population above equilibrium. This value of *σ* serves as a proxy for the strength of the system’s resistance to the disturbance introduced by temperature variability: a high value of *Z* signifies continued population growth despite intense temperature noise (a strongly resistant system), while a low value of *Z* reflects a quick collapse of growth rates below equilibrium at even low levels of noise (a sensitive system). Moreover, *Z* is well-defined because populations inevitably die out when sufficiently extreme temperature patterns are experienced, pushing growth factors far below one eventually.

We performed a parameter sweep of *Z* across the natural domain of the parameters (*h, g*). For the deterministic parameters, we take (*h, g*) ∈ [0°C, 30°C] × [0°C, 25°C], as was conducted in (Lewkiewicz et al., 2022). For a conceptualization of the methodology, see Fig. 2. To quantify the population dynamical changes induced at *σ* = *Z* that underlie the collapse of the growth factors, we measured changes in total annual mortality and egg-laying between the baseline case (no noise) and the *σ* = *Z* case. For each *T*_*σ,k*_(*t*), we compute the mortality fraction, 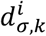, which represents the fraction of the life stage *i* population that dies over the course of the simulation. We then compute the median of the 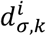 at *σ* = *Z*, denoting it by 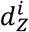. Letting *d*^*i*^ represent 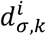 with no temperature variability (*T*(*t*) = *D*(*t*)), the quantity 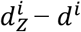 measures how significantly mortality in life stage *i* has changed from its baseline with the introduction of variability at resistance level *Z*. If 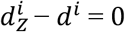, mortality levels are the same at *σ* = *Z* as they are at baseline, and the growth factor collapse is not being driven by changes in mortality in life stage *i*. If 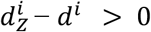, then mortality has increased with temperature variability, and mortality in life stage *i* may be responsible for the collapse of the growth factors. In an analogous manner, we let *o*_*Z*_ denote the median eggs laid per motile at *σ* = *Z*, and let *o* denote the median eggs laid per motile at baseline with no temperature variability, and then consider *o*_*Z*_ – *o*. All growth factor changes result from either excess mortality or a decrease in egg-laying (or both), and the quantities 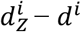 and *o*_*Z*_ – *o* allow us to isolate the contributing factors to population resistance and sensitivity across the (*h, g*) parameter domain.

### 2.4 U.S. Mean Temperature Data

Mean temperature profiles at specific U.S. locations and across the continental U.S. are referred to throughout the remainder of the manuscript as part of the analysis (see Figs. 2, 6, 8, and 9). The average annual mean temperature and average annual temperature amplitude at each location were fitted to the sinusoidal function of Eq. (7) using harmonic regression. To obtain the data used in the fitting, average day-of-year temperatures were calculated from SCDNA data as the average of the recorded temperatures at each location for each day of year from 2008 to 2018 (Tang et al., 2020). The resulting data set was then interpolated on a 0.25-degree grid covering the continental U.S., and the sinusoidal approximation was calculated at each grid point. The boundary of the continental U.S. in (*h, g*) parameter space is the (approximate) boundary of the set of (*h, g*) values across the grid (see Figs. 6 and 8). The approximate (*h, g*) pairing for a particular US location, such as the cities marked in Fig. 8, is the (*h, g*) pairing of the grid point closest to that location.

## 3. Results

### 3.1 Vital Rate Effects on Diapausing and Non-diapausing R_0_ Patterns

In order to understand the population resistance of the spotted lanternfly (SLF) to climate variability, we began by exploring the influence of the vital-rate functions (Fig. 1) on the annual reproductive number, *R*_0_, across parameter sweeps with respect to annual mean temperature (*h*) and annual temperature amplitude (*g*). It was clear in both the diapausing (Fig. 3a) and non-diapausing (Fig. 3c) versions of our model (Eqs. (1-6)) that these vital-rate functions strongly influence realized population growth rates. Specifically, these functions partition the mean-amplitude temperature parameter space into regions in which temperature profiles cross thresholds of activation of different physiological processes (see Figs. 3b, d), per the model calibration and vital rate functions (Table 1, Fig. 1). Thus, any variability in temperature that pushes a population across region boundaries should strongly affect population growth. Below, we provide a thorough description of the results for the *R*_0_ parameter sweeps that led us to this conclusion.

**Fig. 3:**
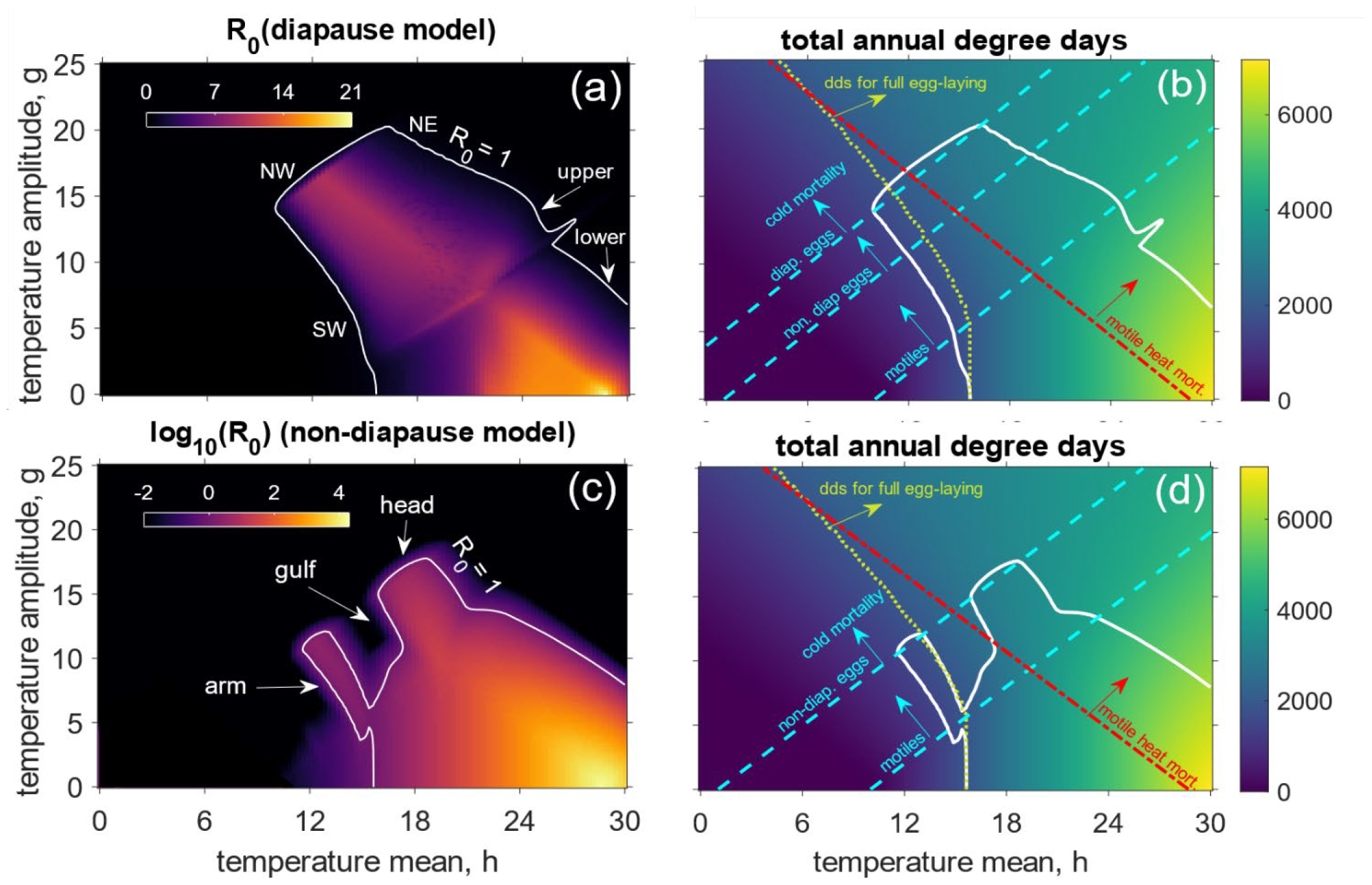
Vital-rate thresholds determine the growth and establishment of diapausing and non-diapausing spotted lanternfly (Lycorma delicatula) populations. (a): Population growth factor (R_0_) across the (h,g) parameter space in the diapause model case with annotations used in the main text denoting the upper and lower regions and the northeast (NE), northwest (NW), and southwest (SW) boundaries of the R_0_ > 1 region. (b): For the diapause model version, total annual degree day accumulation across the (h,g) parameter space, superimposed with the R_0_ = 1 level curve (solid white curve) and threshold lines derived from the vital rate functions denoting regions of activity of vital processes (Fig. 1). Dashed cyan lines denote boundaries above which cold-induced mortality occurs in the labeled population (top line: diapause-lineage eggs, middle line: non-diapause eggs, bottom line: motiles). Below each line, cold does not induce mortality in the associated life stage. The red dash-dotted line denotes the boundary above which heat-induced mortality occurs in motiles. Below this line, heat mortality is not experienced by motiles. The yellow dotted curve denotes the boundary between the part of the parameter space in which the total annual degree day accumulation exceeds the amount needed for an individual to complete the life cycle (right of the curve) and the part in which annual degree day accumulation falls short (left of the curve). Mathematically, this is the level curve DD = 1868.7 degree days. (c): R_0_ in the non-diapause model case (note the log scale) with annotations used in the main text denoting the arm, head and gulf regions. (d): same as (b) for the non-diapause model case.

To understand the effects of the vital-rate functions and boundaries, we produced heat maps of the total number of degree days accumulated in a one-year period (see section 2.1) under the temperature profile of Eq. (7), with the *R*_0_ = 1 level curves from Fig. 3a and Fig. 3c reproduced in Fig. 3b and Fig. 3d, respectively (solid white curves). Also depicted in Fig. 3b and Fig. 3d are the threshold lines. The three parallel dashed cyan lines in each figure have equations of the form *h* – *g* = *b*, thus representing the line of (*h, g*) pairings that produce a temperature profile with an annual minimum of *b* (°C). For the topmost line, for instance, *b* = –3.96°C, which is the threshold temperature below which some fraction of diapause-lineage eggs is killed by cold. In the region above this line, temperature profiles dip below this threshold near the winter minimum, inducing mortality in any existing diapause-lineage eggs. Below the line, temperatures stay above –3.96°C and diapause-lineage eggs do not experience death induced by cold. The middle line, with *b* = 1.04°C, and the bottom line, with *b* = 10°C, analogously split the parameter domain according to the thresholds for non-diapause egg and motile cold mortality, respectively. The dash-dotted red curve has equation *h* + *g* = 28.7°C; above this line, temperature profiles have a maximum annual temperature that exceeds 28.7°C, above which motiles experience heat-induced mortality. The dotted yellow curve in each figure refers to the heat map of total degree day accumulation itself: to the right of this curve, temperature profiles yield an annual degree day accumulation greater than that necessary to drive individual maturation from the moment development commences in an egg through hatching, the juvenile stages, and the completion of the egg-laying stage (here, 1868.7 degree days (Calvin et al., 2023) [D. Calvin personal communication]). To the left of this line, annual degree day accumulation falls below 1868.7 degree days and individuals are not able to reach full egg-laying capacity within the span of one year.

The *R*_0_ parameter sweeps for the diapause and non-diapause model versions were strongly affected by interactions among the vital-rate functions (Fig. 1). In the diapause model version (i.e., the version that best reproduces the diapause possible in the U.S. invaded range), the parameter space splits into three regions with qualitatively distinct *R*_0_ patterns (Fig. 3a). In the region exterior to the *R*_0_ = 1 level curve (black area), *R*_0_ is less than 1; here, temperature profiles guarantee either high rates of cold- or heat-induced mortality or degree day accumulation that is too low for egg-laying sufficient to maintain a population (or both). The *R*_0_ = 1 curve surrounds two qualitatively distinct regions, for both of which *R*_0_ > 1: the centrally located moderate-mean/moderate-amplitude “upper” region, and, to its lower right, the high-mean/low-amplitude “lower” region (labeled in Fig. 3a). In the upper region, *R*_0_ has a maximum of ∼10, which is attained along the motile heat death line (*h* + *g* = 28.7°C) described above (red dashed line in Fig. 3b). For temperature profiles above this line, motiles experience heat death in the summer—potentially before and/or during egg-laying—with greater duration and severity the farther above the line a temperature profile lies. For *h* and/or *g* values far above this line, high summer mortality rates lead to limited fall egg-laying, an effect that becomes severe enough at the northeast boundary of the upper region (labeled NE in Fig. 3a) for *R*_0_ to fall below 1. (Note: Throughout the remainder of the manuscript, the labeled segments of the boundary of the upper region in Fig. 3a will be referred to by the acronyms “NE,” “NW,” and “SW,” which, of course, refer to their ordinal direction placements. When using cardinal and ordinal descriptions to refer to broad regions of the continental United States, these words will be written out in full (e.g., “the northeast” refers to the US region containing New England and neighboring states). Below the motile heat death line, lower annual means and amplitudes correspond to lower egg-laying rates due to slower and generally diminished degree day accumulation. As *h* and/or *g* decrease below the line, summer motile development and egg-laying continue to slow, further reducing *R*_0_; eventually, the threshold curve for full annual degree day accumulation is crossed (yellow dotted curve in Fig. 3b) and the SW boundary of the upper region is reached, indicating that degree day accumulation has fallen too low for population persistence (SW in Fig. 3a). From the approximate symmetry of this *R*_0_ pattern across the motile heat death line (see the *R*_0_ ridge in Fig. 3a that corresponds to dashed red line in Fig. 3b), we infer that the *R*_0_ value at a point (*h, g*) is correlated primarily with its proximity to the motile heat death line. Indeed, *R*_0_ values are essentially unrelated to proximity to the diapause egg death line (top dashed cyan curve in Fig. 3b), except in the small sliver of the upper region that lies above this line, where winter egg death quickly brings *R*_0_ below 1. In the lower region, *R*_0_ attains its absolute maximum value of ∼21 at (*h, g*) ∼ (28.7°C, 0°C). This temperature profile (generally unrealistic outside of tropical areas) optimizes *R*_0_ because 28.7°C is the highest temperature at which populations are not subjected to any form of temperature-induced death, allowing for maximal development speeds with no excess mortality (see the brightest point in Fig. 3a and its corresponding location in Fig. 3b). Under this temperature profile, temperature-independent background death is the only source of mortality in any population: a filter built into the egg-laying rate removes 40% of all eggs laid, while motiles perish at a rate of 0.012 deaths per capita per day. Without this background motile death, one would expect an *R*_0_ of 30 (that is, 60% of 50, a female’s maximum lifetime egg-laying capacity). However, this number is reduced to ∼ 21 due to the background motile death that eliminates motiles before and throughout the egg-laying process.

As in the upper region, the motile heat death line (*h* + *g* = 28.7°C) creates a sharp gradient at which *R*_0_ drops rapidly due to the introduction of summer motile heat mortality. In the wedge-shaped subset of the lower region that lies below this line, populations experience no excess death due to temperature; performance is instead limited by slower degree day accumulation interacting with basal mortality. Temperature profiles in this region are confined entirely to the interval (*T*_dev_, *T*_plat_) on which daily degree day accumulation is an affine linear function of *T*(*t*) (see Fig. 1a), implying that total annual degree day accumulation, 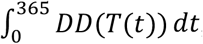, depends only on the annual mean, *h*, of the profile *T*(*t*). With *DD*(*T*(*t*)) = *xT*(*t*) + *d* in the interval (*T*_dev_, *T*_plat_), for some constants *x* and *d* from the data-calibrated degree day function (Smyers et al., 2021), this can be seen by computing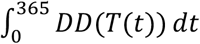 explicitly as such:

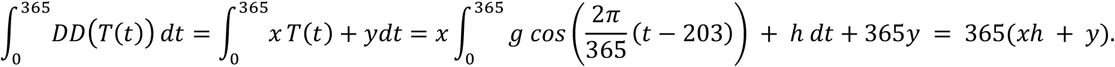

The amplitude (*g*) only affects the trajectory of cumulative degree day accumulation over the course of the year—slowing it in the winter and hastening it in the summer—which may interact with basal mortality rates to induce small variation in *R*_0_ along the vertical lines of constant mean (*h*) in the parameter domain. However, as is apparent in Fig. 3a, *R*_0_ is correlated primarily with the annual mean temperature, decreasing as *h* does until the threshold for full degree day accumulation (yellow dotted curve) is crossed, followed quickly by the *R*_0_ = 1 curve. In summary, population-dynamical performance for the diapause model version under mean temperatures (*R*_0_) is shaped primarily by motiles’ experiences of temperature patterns. Strong performance is experienced at temperatures along the motile heat death line, while there is diminishing performance below the line (at lower amplitudes) due to insufficient degree day accumulation and also above the line (at higher amplitudes) due to summer motile death. The only element of the *R*_0_ > 1 region shaped primarily by egg mortality is the position of the NW boundary of this region (Fig. 3a), which is determined by the line of cold-induced mortality in diapause-lineage eggs.

Maximal achievable growth in the non-diapause model version (i.e., the version that may reproduce dynamics observed once SLF invades more southern and western areas of the U.S.) is orders of magnitude larger than in the diapause model version, but the range of establishment in the parameter domain is smaller (note the log scale and difference in the shaded area in Fig. 3c vs. Fig. 3a). The upper extent of the non-diapause *R*_0_ > 1 region (Fig. 3c) is determined by the threshold for cold-induced mortality in non-diapause eggs (upper dashed cyan curve, Fig. 3d). The higher threshold for cold mortality in non-diapause eggs thus eliminates establishment potential in much of the upper region of the diapause model case (compare shaded area in Fig. 3c with Fig. 3d). Although not visually apparent from the type of sharp gradients in the *R*_0_ pattern seen near or along lines representing biological thresholds in the diapause case, the motile cold mortality line (lowest dashed cyan curve in Fig. 3d) corresponds to a fundamental divide between contrasting phenologies and *R*_0_ behavior in the non-diapause case. Above this line, motiles do experience cold-induced death that can limit egg-laying, leading to lower *R*_0_ values. Moreover, the arrangement of this region, consisting of two distinct areas with *R*_0_ > 1 (labeled “arm” and “head” in Fig. 3c) separated by a gulf in which *R*_0_ < 1, results from temporal interactions between varying motile development rates and the onset of cold mortality (see crossing threshold lines in Fig. 3d). Across this region, a roughly age-synchronized motile population emerges after the summer temperature peak, which lays its own eggs in the fall as development slows and the period of cold-induced mortality approaches; it is the fate of this second cohort of eggs that determines *R*_0_. In the “arm” region, where *h* is generally lower, development is slow enough to prevent a sufficient number of second-cohort eggs from hatching into motiles before the motile cold mortality threshold is crossed, killing off the motile population (Appendix 1: Point 1, no noise). The eggs, however, survive the cold because this entire region lives below the non-diapause egg cold mortality line. In the gulf, development is fast enough to force complete hatching of all second-cohort eggs, but too slow for them to lay another cohort before they are killed by the cold (Appendix 1: Point 2). In the “head” region, development is now fast enough to allow the second-cohort eggs to lay a third cohort that survives the winter, bringing *R*_0_ back above 1. Behavior in this region illustrates the utility of diapause in preventing premature hatching that transitions individuals from a life stage less vulnerable to cold to one more vulnerable. Below the motile cold mortality line, there is a year-round presence of both eggs and motiles that supports constant, rapid flow through the lifecycle and explosive population growth. In this region, populations experiencing temperature profiles above the motile heat death line (resp., below the line) are still limited by heat-induced mortality (resp., slower development), as in the diapause model case.

### 3.2 Population Resistance to Climatic Variability

We developed a metric of population resistance, *Z*, to quantify how annual growth factors depart from the baseline *R*_0_ as temperature variability intensifies. Specifically, the metric is the standard deviation of the noise term *σ* (Eq. (8)) that eventually drives growth factors below 1 (Eqs. (11-12)). To understand the relationship between *Z* and *R*_0_ we performed a parameter sweep across *h* and *g*, as we did for *R*_0_ (Fig. 3) and plotted Z for the diapause and non-diapause model versions with several *R*_0_ level curves superimposed on the *Z*-heat map (Fig. 4). We also estimated the changes in egg and motile mortality and egg-laying between baseline average temperature conditions and resistance-level noisy conditions (Fig. 5).

**Fig. 4:**
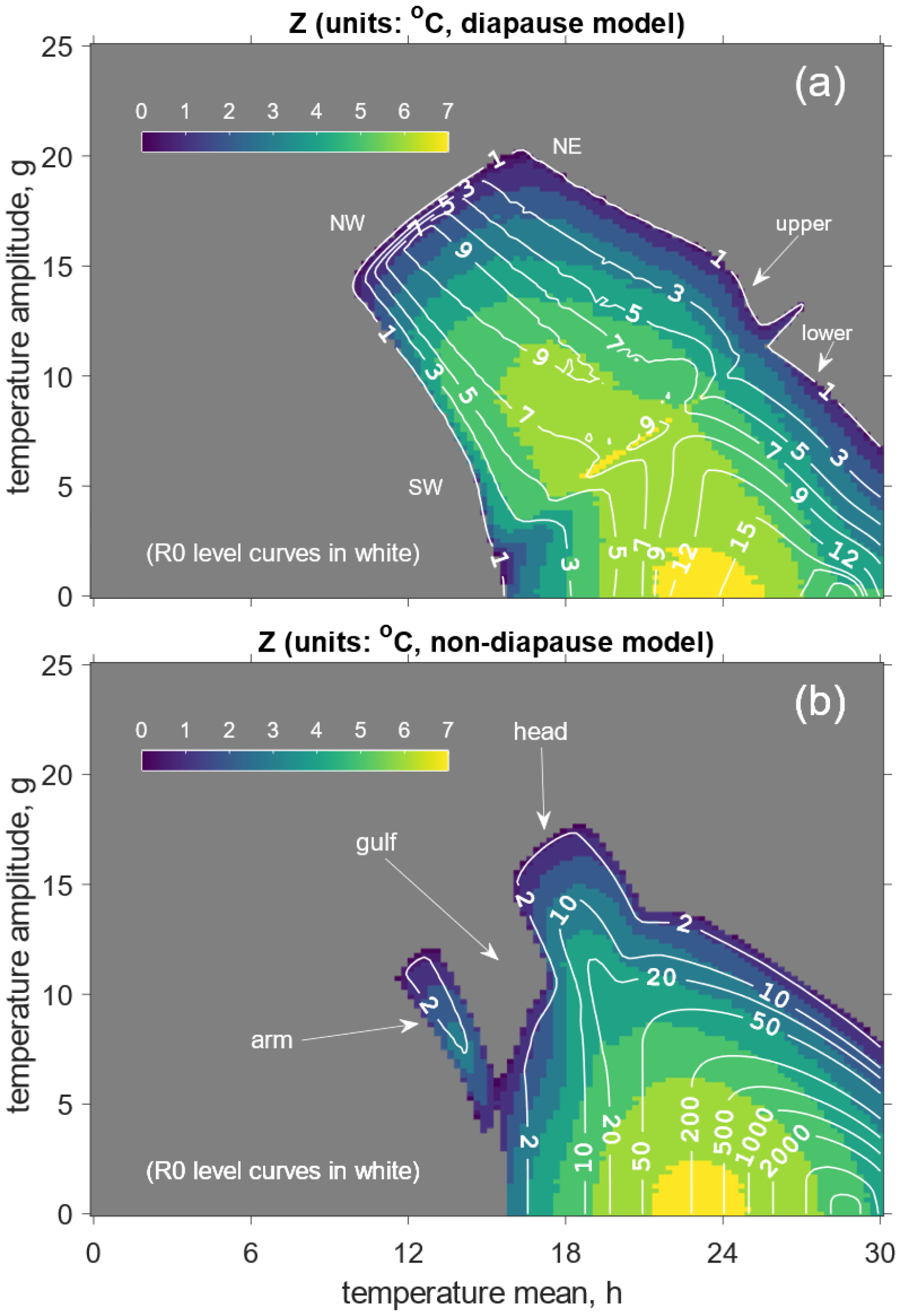
Population resistance to climatic variability (Z) and population growth factor (R_0_) are complementary establishment metrics. The metrics are complementary because the R_0_ level curves (white curves with numbers indicating R_0_ value derived from Fig. 3a) are not tangent to Z level sets for either the diapause and non-diapause model variations. (a): Heat map of Z in the diapause model variation. Gray covers the area in which R_0_ < 1, where Z is not defined (Fig. 3a). (b) Heat map of Z in the non-diapause model variation.

**Fig. 5:**
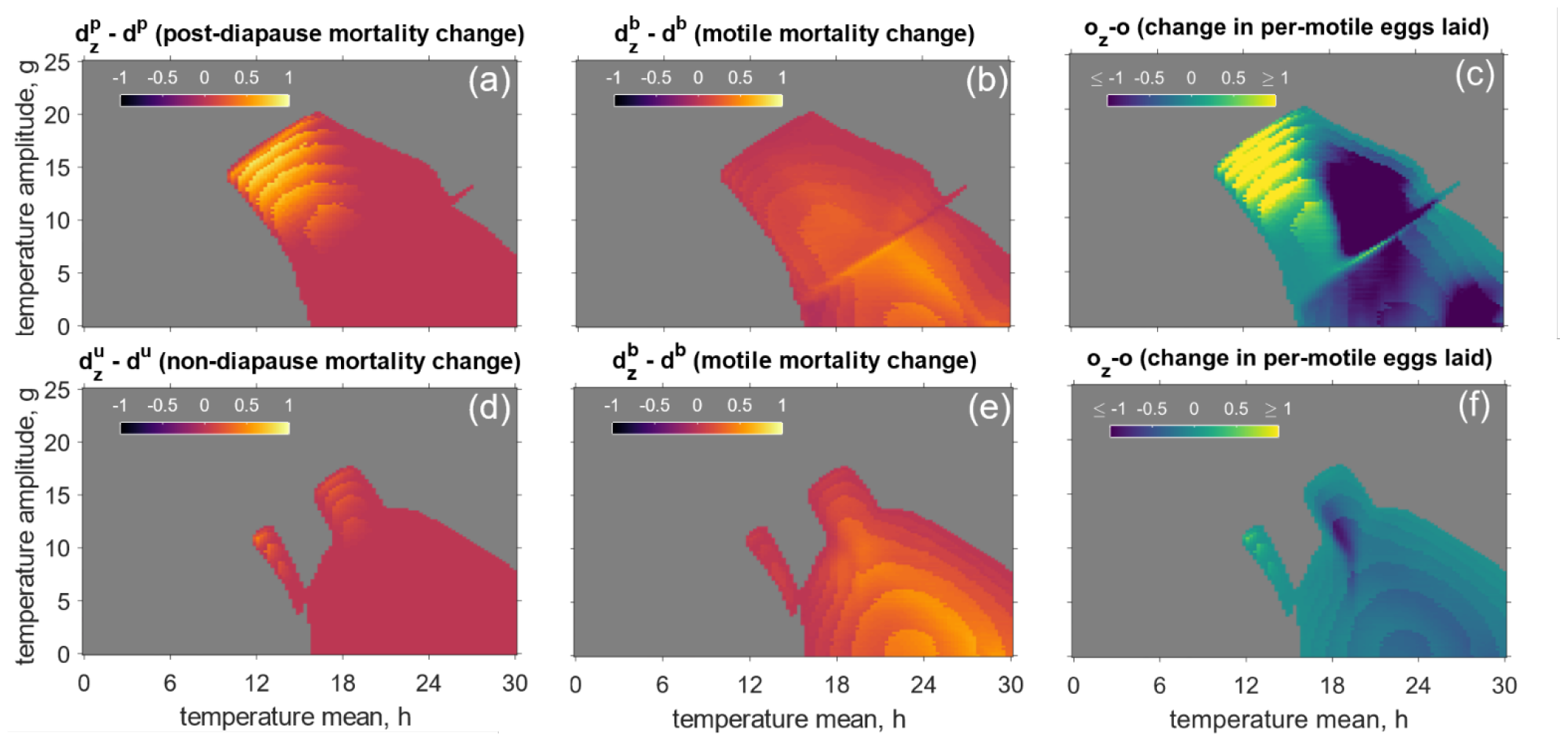
Annual temperature means and amplitudes determine how climatic variability affects population growth via changes to mortality and fertility. Each panel represents change in post-diapause egg mortality, motile mortality, and per-motile eggs laid (fertility) between the baseline (no noise) and resistance level Z in the diapause model version (a-c) and non-diapause model version (d-f). In (a,b,d,e), a value of +1 means that no individuals in the relevant life stage die at baseline but all die when stochastic noise variability σ = Z, while a value of -1 means that all individuals in the relevant life stage die at baseline but none die when σ = Z. In (c,f), o_z_ – o can take any value in the real numbers; a positive number indicates that noise contributes to higher per-motile egg-laying, whereas a negative number indicates that noise contributes to lower per-motile egg-laying. The higher the magnitude of o_z_ – o, the more significant the change.

In the diapause model case, *Z* achieves its maximum value in a small subset of the lower area in which *h* ≈ 21 – 25°C and *g* ≈ 0 – 2°C (Fig. 4a). This value is achieved at slightly lower mean temperatures than that at which *R*_0_ is optimized ((*h, g*) = (28.7°C, 0°C), Fig. 3a). Indeed, temperature profiles that effectively resist the effects of noise are, in general, ones in which the annual range remains between—and well separated from—the threshold temperatures for cold and heat mortality but are also as warm as possible to support quick development and high rates of egg-laying. Resistance is not highest for the temperature profile defined by (*h, g*) = (28.7°C, 0°C) because this point lies on the motile heat mortality line itself, where any noise at all will periodically induce motile death (compare Fig. 4a to Fig. 3b). The temperature patterns with means a few degrees lower balance the maximization of distance to mortality thresholds with the maximization of warmth that supports development and egg-laying. Resistance decreases outward from its optimal region (bright yellow in Fig. 4a) as the boundary of the *R*_0_ > 1 region is approached; populations are naturally less resistant on the boundary, where establishment potential is low under mean conditions and noise typically exacerbates that poor baseline performance. While the decline is gradual near most of the *R*_0_ = 1 boundary, the gradient in Z values is notably sharp along the SW boundary of the upper region (Fig. 4a).

In the upper region of the diapause version of the SLF model, Z is determined jointly by proximity to the diapause egg cold mortality threshold line and the motile heat mortality threshold line. (Fig. 3b, 4a). This is due to the fact that in the upper region of the *R*_0_ parameter domain, populations are univoltine, with all eggs entering diapause (Appendix 1: Point 3, no noise). The generally higher amplitudes (*g*) in the upper region support rapid development and a brief period of intense summer/fall egg-laying, while also placing summer and winter temperature extrema near or past the thresholds for excess mortality. The *Z* and *R*_0_ level curves are nearly perpendicular to each other below the motile heat mortality line and nearly parallel above the line. Below the motile heat mortality line (red dashed line in Fig. 3b), resistance runs parallel to the diapause egg cold mortality threshold (see the top, cyan-colored dashed line in Fig. 3b) because noise inhibits populations through cold trends that pull winter temperatures below the diapause egg mortality threshold. This source of mortality is clear in Fig. 5a, which indicates that post-diapause egg mortality increases significantly at resistance level *Z* relative to its baseline in this upper region where *h* ≤ 18°C and *g* ≥ 8°C. Interestingly, per-capita egg-laying is also higher in this region when noise is introduced (Fig. 5c), indicating that while noise raises egg-laying rates, these gains in egg-laying cannot compensate for diapause egg death. Above the motile heat mortality line (Fig. 3b), resistance runs parallel to the NE boundary, as warm trends in the summer months exacerbate existing heat-induced motile mortality that diminishes egg-laying to bring populations below equilibrium. A severe decrease in per-motile egg-laying occurs in this *h* ≥ 18°C part of the upper region (Fig. 5c). Along the SW boundary, *R*_0_ is limited by insufficient egg-laying (Fig. 3a,c), but the presence of noise does not appear to lower population persistence by lessening egg-laying near that boundary the way it does near the NW and NE boundaries by increasing egg and motile mortality (Fig. 4a). This reflects the dichotomy between how temperature noise interacts with development and egg-laying through the degree day function (Fig. 1a) and how it interacts with temperature-induced death through the mortality curves (Fig. 1c). Noise induces (often sporadic) bouts of mortality as it draws temperatures back and forth across mortality thresholds, while any warm (resp., cold) trend in noise increases (resp., decreases) egg-laying. Far from temperature mortality thresholds, then, median egg-laying behavior remains stable in a noisy environment, rendering it the less influential factor in resistance outcomes. In the lowest amplitude part of the upper region (*h* ≤ 18°C, *g* ≤ 8°C), egg-laying rates are low at baseline and vary significantly with noise, while noise may push temperatures below the motile mortality threshold in the fall and spring to either interrupt egg-laying or kill newly hatched motiles. At resistance level *Z*, the responsibility for population collapse is shared between these processes and diapause egg death in the winter, which explains the lack of dominance of any particular aspect of change in (Fig. 5a-c).

In the lower region, right at the transition between the upper and lower regions, the loss of motile cold mortality and overall warmer temperatures cause a fundamental shift in the underlying phenology away from the age-synchronized, univoltine structure characteristic of the upper region (Fig. 3a-b, 4a). In the lower region, diapause egg-laying typically occurs continuously throughout the fall, building up a slow-developing population that hatches the following summer (Appendix 1: Point 4, no noise). The few motiles remaining after the winter solstice lay non-diapause eggs that hatch quickly, initiating cycles of continuous egg-laying through the non-diapause pathway. Summertime then sees a massive influx of juveniles from both lineages. In the lower region, egg mortality is unaffected by the presence of temperature noise because temperature profiles are far above the egg cold mortality thresholds. Noise inhibits population growth across the lower region by increasing motile mortality and decreasing per-motile egg-laying, the latter to varying extents (Fig. 5b-c). Although motiles are often present throughout the year in the lower region, the largest population occurs in the summer when overwintered diapause eggs hatch, roughly in unison (Appendix 1: Point 4, no noise). Noise exacerbates high summer temperatures to kill this newly-hatched motile cohort quickly before it lays any eggs, in turn significantly lowering annual per-motile egg-laying in parts of the lower region.

In the non-diapause model version (Fig. 4b), optimal resistance occurs at temperatures similar to those at which it occurred in the diapause model case (*h* ≈ 21 – 25°C, *g* ≈ 0 – 2°C), with both *Z* and *R*_0_ decreasing approximately radially from their respective optima. Resistance is poor above the motile cold mortality threshold line as noise pushes temperatures below the non-diapause egg cold mortality threshold in the winter, killing eggs (Fig. 5d, see thresholds in Fig. 3d). As in the lower region of the diapause model case, noise decreases populations by increasing motile mortality (Fig. 5e). In this case, however, per-motile egg-laying rates are nearly unchanged at resistance level *Z* in comparison with their baselines (Fig. 5f)—in stark contrast with the diapause model case, in which noise dramatically alters per-motile egg-laying across much of the parameter space (Fig. 5c). This is due to the continuous presence of both eggs and motiles moving through the life cycle in the non-diapause model case, which lessens the overall impact of bursts of mortality induced by noise (Appendix 1: Point 5).

### 3.3 Diapause relative advantage under variable climates

As SLF spread across the U.S., they will encounter warmer and less seasonal climates with increasing frequency. In such climates with higher annual mean temperatures and lower annual mean amplitudes, diapause may no longer be advantageous. Indeed, non-diapausing populations under our model exhibit much higher population growth in warmer and less seasonal regions, suggesting a relative advantage for non-diapausing populations (note the log scale difference between Fig. 3a,c). However, temperature variability may influence this advantage.

Under some variable temperature conditions, non-diapausing populations might have a higher *R*_0_ and lower *Z* than diapausing populations. Under such conditions, non-diapausing SLF populations could emerge in nature. To test how variability influences diapause, we estimated relative advantage with respect to *R*_0_ and *Z* across our annual mean (*h*) and amplitude (*g*) parameter sweep (Fig. 6). For this analysis we denoted by 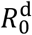 and 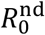 the diapause and non-diapause *R*_0_ values, and analogously for *Z*. In Fig. 6a, we plotted *Z*^nd^ – *Z*^d^, the disparity between Z-values in the diapause and non-diapause cases, which is positive when non-diapause has higher resistance and negative when diapause does. (While the actual range of values spans -5.5 to 2, it is plotted in linear scale from -3 to 3 for visual symmetry.) Non-diapause populations have the *Z*-advantage at very high means and very low amplitudes, while diapause populations have the advantage everywhere else. In Fig. 6b, we plot the quantity 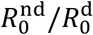 in natural logarithmic scale; values above 0 occur when diapause has a higher reproductive number, and values below 0 occur when non-diapause does. (A logarithmic scale is used here because the range of 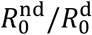 spans many orders of magnitude.) With respect to *R*_0_, non-diapause has the advantage over a significantly larger subset of the pertinent part of parameter space than it did with respect to *Z*.

**Fig. 6:**
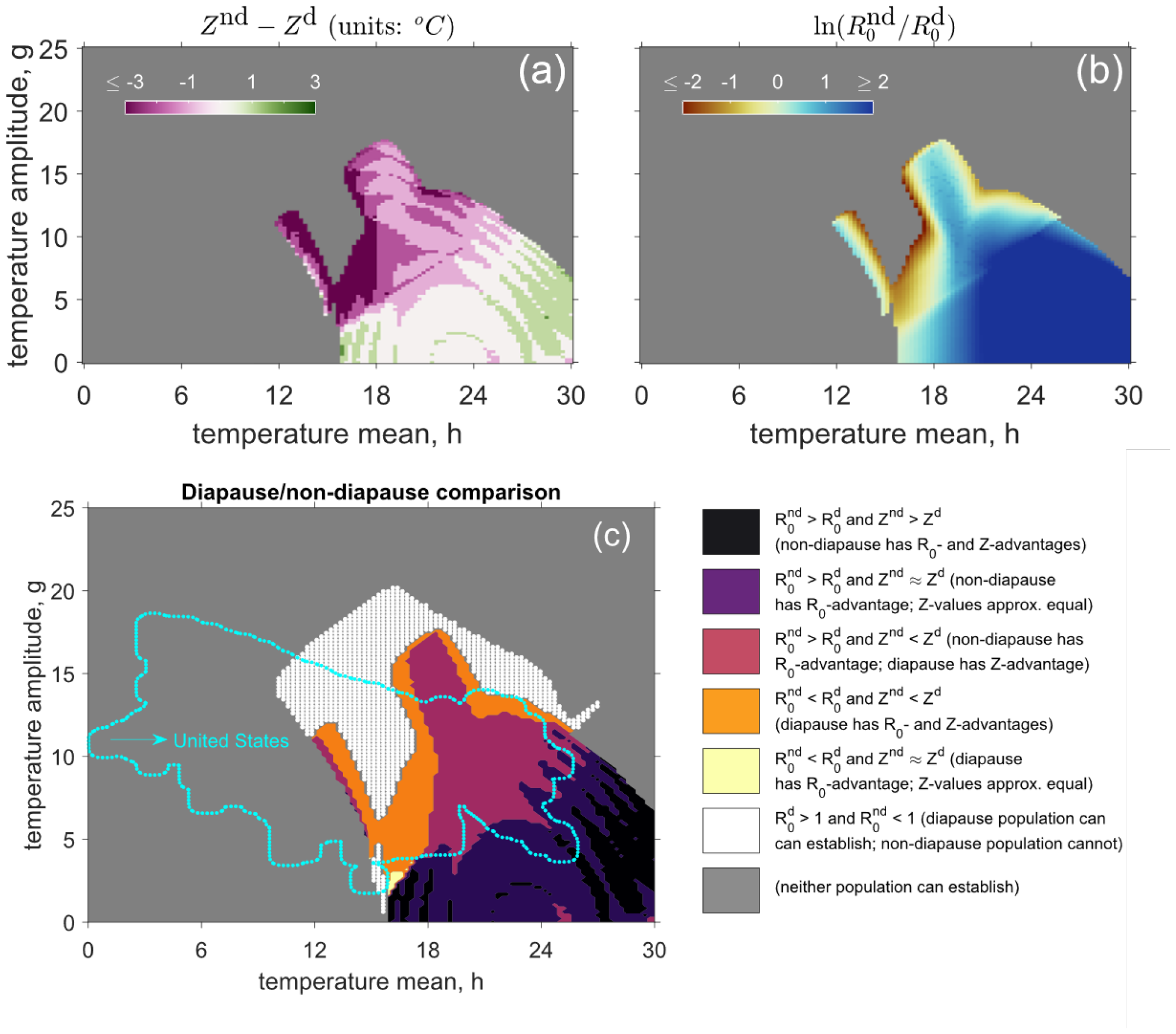
Diapausing populations have a relative advantage due to climatic variability throughout most of the U.S. Across much of the diapause upper region, non-diapause populations cannot establish (white dots). Near the border of the non-diapause establishment region just below it (orange area), diapause populations have a higher R_0_ and Z. Just below that (burgundy region), non-diapause populations gain the R_0_ advantage, so that they outperform diapause populations in terms of baseline reproductive number but still fall short in terms of resistance. Below that, in the lower right corner (black and dark purple areas), non-diapause populations have the advantage in terms of both R_0_ and Z. The outline of the region approximating mean United States temperature profiles is superimposed on the parameter space (dotted cyan curve), showing that diapause populations are more resistant in most of the parts of the continental United States in which establishment is possible (the northeast, south, and much of the Midwest), despite having a smaller reproductive number in warmer areas.

In Fig. 6c, these dominance relationships are summarized categorically across the temperature parameter space. Only in a small area of the (*h,g*) parameter space did non-diapausing populations have both a higher *R*_0_ and a higher (or equivalent) *Z* than diapausing populations (see black and dark purple areas in Fig. 6c). As temperature means decrease and amplitudes increase, diapause becomes the more advantaged population. In the upper part of the upper region (white dots in Fig. 6c), non-diapausing populations cannot establish, while diapausing populations can. Near the boundary of the *R*_0_ > 1 range for non-diapausing populations (orange area), diapause has the advantage of a higher *R*_0_ and *Z*, as one would expect. At slightly lower amplitudes and higher means, though, diapausing populations lose the *R*_0_ advantage, but diapausing populations are still more resistant (burgundy region in Fig. 6c).

These results indicate that the incorporation of climatic variability into the model shows that the presence of variability creates an advantage favoring diapause populations. In almost the entirety of the SLF-hospitable U.S., diapause populations are more resistant than their non-diapause counterparts. We estimated the area of the parameter sweep domain that corresponds to U.S. temperature profiles (bright blue dotted curve outline in Fig. 6c). When climate variability is included, it is the case that in much of this U.S. parameter area either 1) only diapause populations can survive under mean temperature conditions, 2) diapause has both the *R*_0_- and *Z*-advantages, or 3) diapause boasts the *Z*-advantage and non-diapause the *R*_0_-advantage. In the third case, although the non-diapause *R*_0_ value may be significantly larger than the diapause *R*_0_ value for some temperature profiles, a diapause population’s realized growth rate under stochastic temperature conditions is very likely to be higher if the noise level exceeds *Z*^*nd*^ and falls short of *Z*^*d*^. If the noise level also falls short of *Z*^*nd*^, diapause and non-diapause populations would both most likely grow, but non-diapause populations might experience a greater lessening of growth factors due to the fact that noise generally impacts a non-diapause population more severely under these circumstances. A small portion of the U.S. temperature region—representing southern Florida—overlaps with the area of non-diapause advantages in both *R*_0_ and Z (Fig. 6c, dark purple area within dotted cyan curve). Only this small part of the country exhibits climates where natural selection may be relaxed enough to allow non-diapausing populations to proliferate (see the Discussion).

### 3.4 Can noise raise growth rates?

Having identified the noise levels required to suppress populations below equilibrium across the deterministic temperature parameter space, we provide an example of climatic variability raising growth rates. This endeavor is prompted by the presence of areas within the *R*_0_ > 1 regions in which egg-laying rates rise in the presence of noise: the lower-mean subset of the diapause-model upper region (Fig. 5c, *h* ≤ 18°C, *g* ≥ 8°C) and the upper arm region of the non-diapause model (Fig. 5f). While both of these regions lie in close proximity to egg cold mortality lines (Figs. 3b,d) and thus present lower resistance levels, the apparent potential for high egg-laying rates suggests that sub-resistance-level noise may often raise growth factors. To probe this possibility, we investigate trends in the range of the growth factors as the standard deviation parameter, *σ*, increases at a pair of points near the overlap between the diapause and non-diapause high egg-laying regions (Fig. 7). We plot the 300 values of the growth factor, *G*_*σ,k*_ (Eq. (11)), computed for each *σ* = 0.1, 0.5, 1, 2, …, 8 at the points (*h, g*) = (10.8, 13.2) (Point 6, Fig. 7b) in the diapause case and (*h, g*) = (12.4, 10.7) (Point 1) in the diapause case (Fig. 7c) and the non-diapause case (Fig. 7d). These two points, 1 and 6, exhibit average temperature profiles found in the western U.S., including California.

**Fig. 7:**
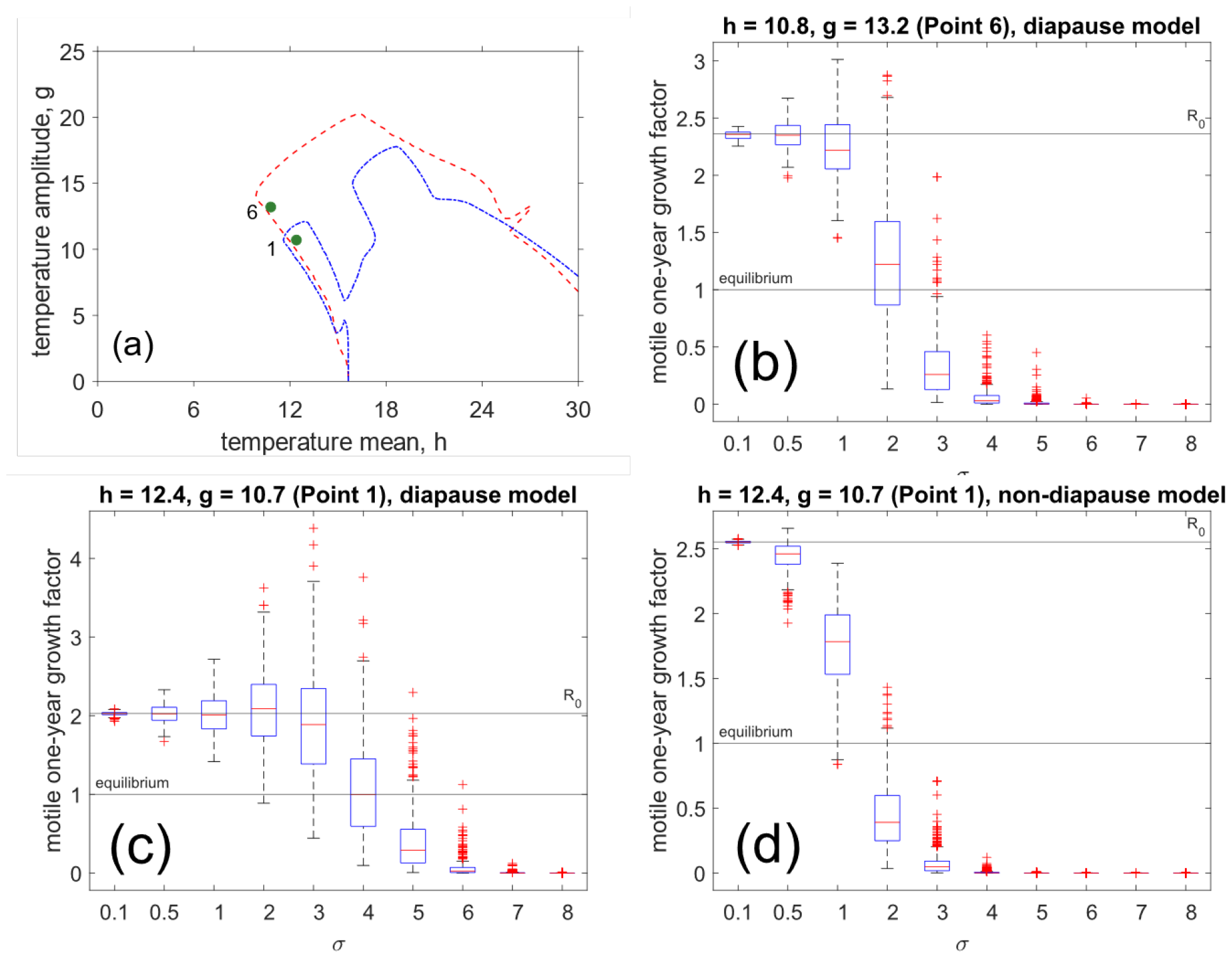
Climatic variability can at times greatly increase spotted lanternfly population growth by increasing fertility. Box plots of growth factors for increasing σ for two deterministic temperature profiles in the low-egg-laying portion of the (h, g) parameter domain that correspond to climates found in the Western U.S. (Fig. 8-9). (a): The locations of points 6 and 1 in the (h, g) domain with respect to the R_0_ = 1 level curves in the diapause and non-diapause model cases (dashed red curve and dash-dotted blue curve, respectively). (b): box plots for Point 6 in the diapause model case, in which growth rates are limited by the close proximity of the temperature profile to the diapause egg mortality threshold line (Appendix 1: Point 6). (c): box plots for Point 1 in the diapause model case, in which higher annual temperatures support growth well beyond R_0_ through σ = 3 (Appendix 1: Point 1, diapause). (d): box plots for Point 1 in the non-diapause model case, in which R_0_ is an upper bound on potential growth factors due to warm temperatures and the absence of diapause promoting quick movement through the egg-to-motile pathway and subsequent population collapse at the onset of winter egg and motile death (Appendix 1: Point 1, non-diapause).

Of the two studied points in our parameter sweep, point 6 is only relevant in the diapause model version (it is outside of the positive growth region in the non-diapause model version) and has a minimum winter temperature of –2.4°C, which is 1.56°C above the diapause egg death threshold. In this example, *R*_0_ ≈ 2.36, and heightened egg-laying does expand growth rates up to ≈ 3 in some of the simulations. This level of growth-factor expansion is more modest than that observed at Point 1 (to be described below) due to the proximity of the minimum winter temperature to the diapause egg death threshold. Point 1 has a slightly higher mean and lower amplitude than Point 6 and is within the *R*_0_ > 1 region of both the diapause and non-diapause models. Due to the generally warmer temperature conditions and greater disparity between the winter minimum (1.7°C) and the diapause egg death threshold (–3.96°C), noise boosts egg-laying rates enough to raise growth factors to nearly twice *R*_0_ in the diapause model case (*R*_0_ ≈ 2.03 and growth factors are as high as ≈ 3.6 at *σ* = 3, Fig. 7c). However, at the same point (1) in the non-diapause case, growth factors barely expand beyond the *R*_0_ value of 2.55, rendering *R*_0_ not only an average measure of growth but also an upper bound on it (Fig. 7d).

Temperature variability can, under certain circumstances, greatly increase population growth rates. While noise raises egg-laying rates in the exact same fashion under the diapause and non-diapause models, it simultaneously raises development rates, as egg-laying and development are directly proportional to one another (see description of *E*(*t*) in section 2.1). Thus, the same noise that intensifies egg-laying in the non-diapause model also accelerates development of those eggs to early hatching, leaving them to perish at the onset of winter’s cold. While we provide only one example (Fig. 7), it is a pertinent one that demonstrates the ability of diapause to interact with temperature noise to promote population growth at significantly higher rates than those predicted under mean temperature conditions that are exhibited in the western U.S.

## 4. Discussion

We performed a comprehensive analysis (Fig. 2) of the effect of climatic variability on population growth (*R*_0_) and U.S. establishment potential for the spotted lanternfly (SLF, *Lycorma delicatula*), a fast-spreading grapevine pest at risk of disrupting the global wine market (Fig. 3, Huron et al., 2022). SLF is currently univoltine in its U.S. invaded range (Nixon et al., 2022). Eggs undergo diapause during the winter and exhibit diapause-induced cold protection, but evidence exists that diapause is a labile trait and eggs do not need to enter diapause to develop and hatch (Urban and Leach, 2023). As SLF spreads further west and south from where it is currently established in the northeastern U.S., it is important to understand how climatic variability can impact both diapausing and non-diapausing populations. For this analysis, we developed a new metric of population resistance to climatic variability (*Z*), which is the level of variability that pushes a growing population into negative growth (Fig. 4). We applied this metric to simulations of a partial differential equation model we developed of how temperature determines the SLF *R*_0_ based on a parameterized set of vital rate functions (Fig. 1, Table 1) that determine *R*_0_ and *Z* (Fig. 3). To estimate establishment potential, annual time series profiles for a given location can be decomposed into annual means (*h*) and amplitudes (*g*) that reflect the average temperature trend of the location (Fig. 2). We performed a parameter sweep of *R*_0_ and *Z* in which we incrementally simulated SLF diapausing and non-diapausing populations across a range of *h* and *g* that encompassed average trends found within the continental U.S.

Although the population resistance patterns across the (*h, g*) parameter space are structurally similar in the diapause and non-diapause model versions (Fig. 4), the changes in vital processes that underlie those resistance patterns are noticeably different (Fig. 5). Per-capita egg-laying rates are particularly sensitive to noise in the diapause context (Fig. 5c) in contrast with the non-diapause context (Fig. 5f), and diapause also yields higher egg mortality rates when that process is active (Figs. 5a,d). The effects of noise on non-diapause populational dynamical processes are more subdued and uniform across the parameter space because life stage counts are more temporally constant in this case. This is especially true below the motile cold mortality line (Fig. 3d), where populations grow overall while maintaining a stable egg-motile balance throughout the year (Appendix 1: Point 5). In this situation, having individuals spread out across the SLF life cycle at all times buffers the population against sources of quick annihilation that target a specific life stage. Diapause, however, tends to synchronize populations developmentally and adjust their phenology, potentially eliminating the buffer and sometimes eliciting dramatic reactions to temperature patterns. In essence, diapause interacts with temperature noise to drastically alter vital processes through its reshaping of phenology, with outcomes that vary from little effect on growth (see Appendix 1: Point 1, diapause, for example) to rapid population collapse (Appendix 1: Point 4, noisy). Despite the potential for diapause to, for instance, align SLF phenology disadvantageously with dangerous temperature extremes, our findings indicate that diapause populations should experience higher levels of resistance across almost the entire continental United States (Fig. 6).

In addition to offering insight into the relative performances of diapause and non-diapause populations, the *R*_0_ and *Z* parameter sweeps provide absolute measures of establishment potential. The introduction of *Z*, and its pairing with *R*_0_, forges a link between average behavior and the wider spectrum of possible outcomes that might emerge in the presence of temperature anomalies, such as cold snaps and heat waves, and general stochastic fluctuations (Lyon et al., 2019). This type of analysis is particularly important in assessing establishment potential in highly variable climates, where realized behavior may deviate significantly from mean behavior (Keena et al., 2023). This can be done for any real location of interest by parameterizing the temperature trend (the parameters (*h, g*)) and then extracting the associated *R*_0_ and *Z* values. If temperature noise levels remain far below *Z* in that location, *R*_0_ is likely a robust approximation of the expected growth rate. If, however, noise levels exceed *Z*, populations are more likely to collapse than to establish, regardless of the value of *R*_0_ itself. For reference, a table of (*R*_0_, *Z*) values at ∼ 30,000 U.S. locations in the diapause and non-diapause cases is provided in Appendix 2 (“US Cities Database | Simplemaps.com,” 2023)

For much of the U.S., when resistance to climatic variability is included, diapausing populations may have more potential to establish than non-diapausing populations (Fig. 6). Areas that fall outside of the diapause model’s *R*_0_ > 1 area include the northern Midwest, Rocky Mountains, and Pacific Northwest (Fig. 8). Parts of the U.S. that fall inside the *R*_0_ > 1 region include much of the Midwest, mid-Atlantic, southern California, and the south. Most of these areas (except the hottest parts of the U.S. southeast and southwest) fall in the upper region below the motile heat mortality line (Fig. 8). Parts of the southeast and southwest fall above this line and/or in the lower region. Resistance is high in much of the U.S. south and southern California, with lower resistance in parts of the Midwest and mid-Atlantic. Thus, disregarding other factors not modeled here like precipitation and water relations, there are places within the U.S. where the growth rate and resistance to climatic variability are both high and temperature-based establishment and persistence of the pest if/when introduced is inevitable.

**Fig 8:**
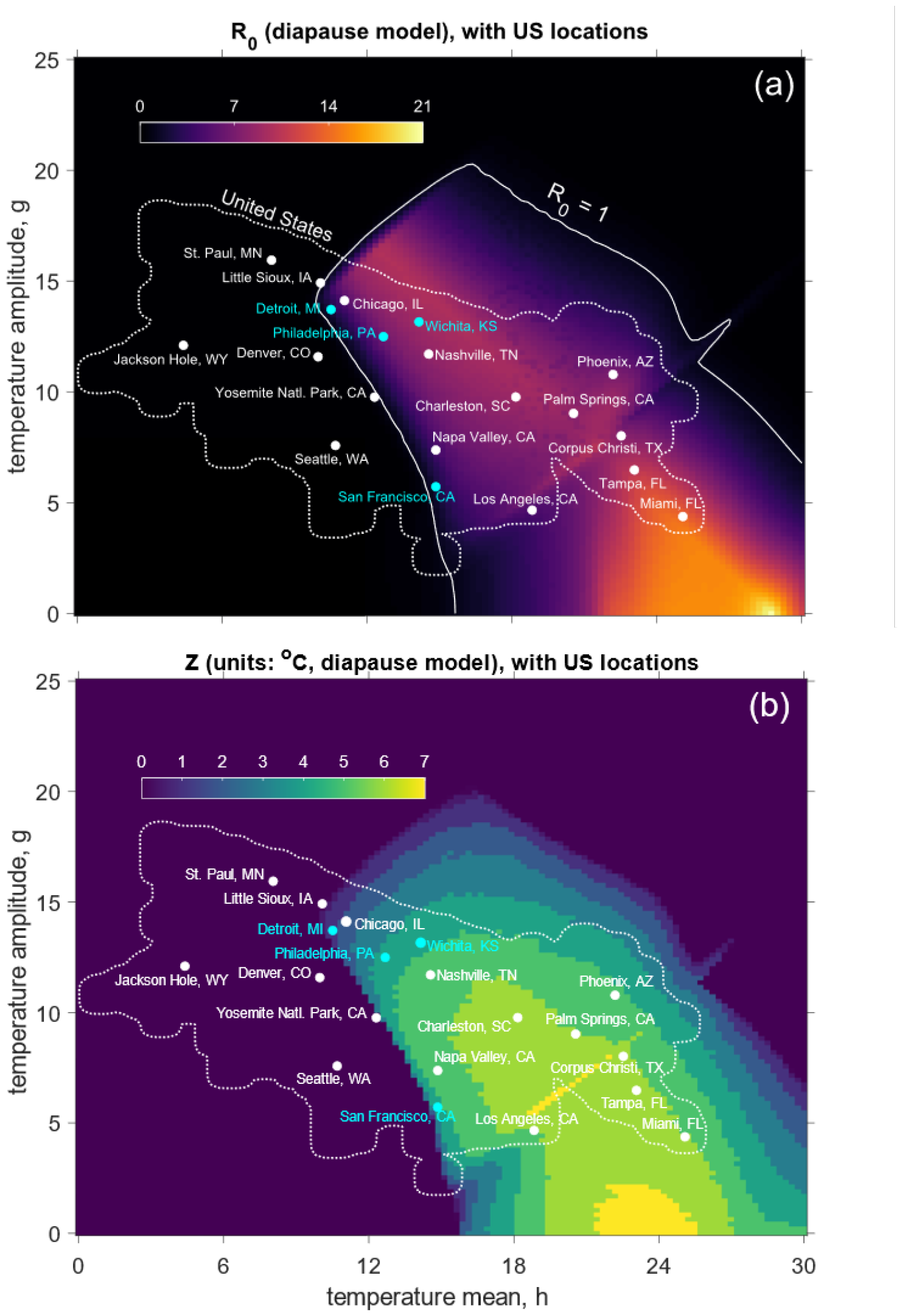
U.S. establishment potential of diapausing populations of spotted lanternfly based on (a) population growth and (b) resistance to climatic variability. Heat maps are R_0_ and Z parameter sweeps over the (h, g) parameter domain (Fig. 3a, 4a), superimposed with the approximate area representing average U.S. temperature trends (interior of dotted white curve) and a selection of U.S. urban areas.

The significance of estimating resistance to climatic variability when determining the establishment potential of an invasive species like SLF is underscored by the fact that the level sets of *R*_0_ and *Z* are generally not tangent to one another (Fig. 4). Indeed, while *R*_0_ and *Z* exhibit a general positive correlation, it is characterized by notable variance, especially in the diapause model case (Fig. 10). This variance stems from the fact that *Z* is a matter of how noise interacts with the deterministic temperature trend to alter its relationship with physiological parameters. A large array of (*h, g*) pairings can produce the same *R*_0_ value while still yielding different realized growth rates under the influence of noise due to differences in the underlying temperature profiles’ relationships with temperature thresholds for physiological processes. Similarly, resistance can be constant across many (*h, g*) profiles with wildly different *R*_0_ values. For example, for diapausing populations, most of the U.S. (Fig. 8) lies in the upper region of the (*h, g*) annual mean and amplitude temperature parameter space below the motile heat death line (Fig. 3b), where *R*_0_ and *Z* are nearly perpendicular (Fig. 4a). For instance, Detroit, MI, and San Francisco, CA, lie on approximately the same *R*_0_ contour line (*R*_0_ = 2.3 and 2.2, respectively), despite their notably different temperature profiles: for Detroit, (*h, g*) = (10.5°C, 13.7°C), whereas for San Francisco, (*h, g*) = (14.8°C, 5.7°C). Detroit’s higher amplitude counterbalances its lower mean, while San Francisco’s higher mean counterbalances its lower amplitude, to lead to similar annual degree day accumulations in both locations (≈ 1.62 × 10^3^ total degree days in Detroit and ≈ 1.69 × 10^3^ in San Francisco) and thus comparable baseline *R*_0_ values. However, Detroit’s winter minimum of –3.2°C lies near the diapause egg death threshold (–3.96°C, Fig. 3b, Table 1), leaving diapausing eggs highly vulnerable to the effects of temperature noise and resulting in a low resistance value of *Z* = 1°C. The annual range of San Francisco’s temperature trend is (9.1°C, 20.5°C), which remains far from all mortality thresholds except that of motiles experiencing cold, resulting in greater stability and a higher resistance value of *Z* = 4°C. The overlap of the temperature profile with motile cold death temperatures is less influential because cold only kills adult motiles for a brief period in the winter after they have laid eggs. In contrast, Philadelphia, PA and Wichita, KS, provide an example of a pairing with the same resistance level, *Z* = 4°C, but different *R*_0_ values (*R*_0_ = 6.3 in Philadelphia, 9.1 in Wichita). Both locations are positioned in the upper part of the upper region and succumb to diapause egg mortality at the same noise level due to their comparable winter minimum temperatures (0.2°C for Philadelphia, 0°C for Wichita). However, Wichita’s higher mean and amplitude boost egg-laying and lead to a 44% higher *R*_0_ (Fig. 8). These examples illustrate the important point that the spectrum of realized outcomes cannot be inferred from average trend predictions alone, and insight in this direction can be derived by considering the proximity of average trends to temperature thresholds, with outcomes quantified by resistance measures such as *Z*.

**Figure 9:**
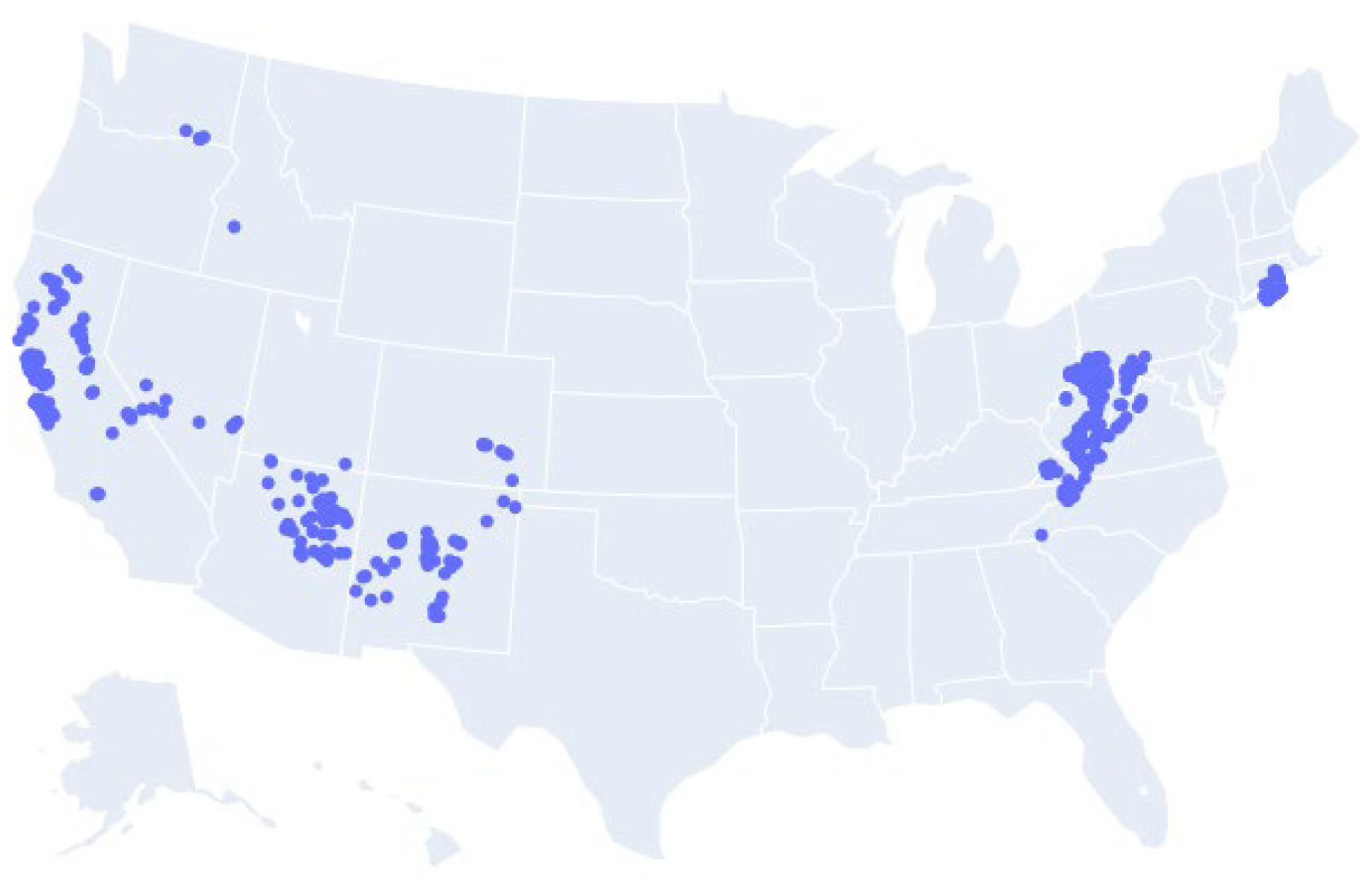
US Locations at which R_0_ may underestimate realized growth: Blue dots indicate cities with temperature trends (that is, best fits of (h, g) to historical average daily mean temperatures) that fall near the southwest boundary of the upper region in the diapause model case and have growth factors that equal or exceed 1.5 x R_0_ at some noise level, σ. Points are deemed “near” the SW boundary see if they fall within the subset of the upper region bounded by the R_0_ = 1 level curve to the left, the curve of full degree day accumulation (yellow dotted curve in Figs. 2(b,d)) to the right, and the diapause egg cold mortality line (topmost dashed cyan curve, Fig. 2(b)) above. Temperature profiles in this region of the (h, g) parameter space are represented across the U.S., including areas across California (notably including Sonoma and Napa Counties, CA), parts of Arizona, New Mexico, Colorado, West Virginia, Pennsylvania, Connecticut, and eastern Long Island.

**Figure 10:**
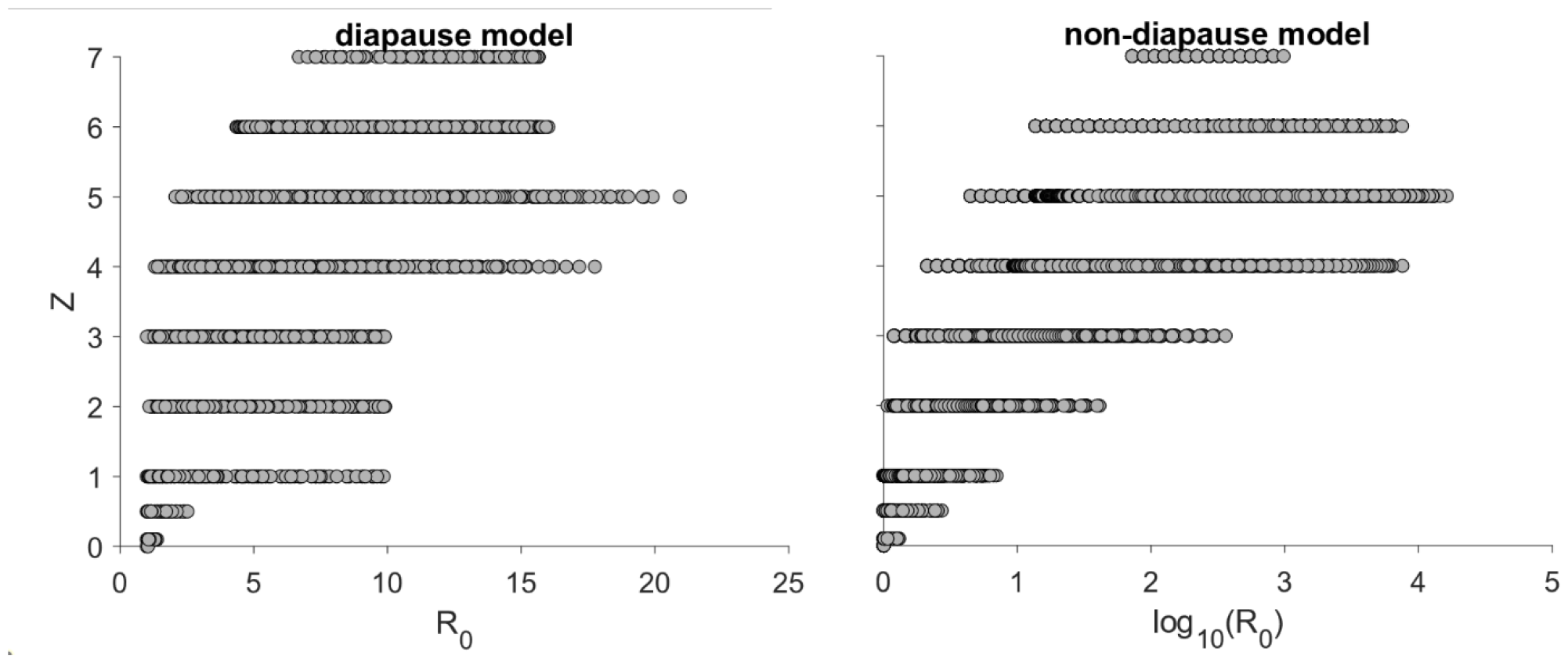
Population growth and climatic-variability resistance are nonsynonymous measures of spotted lanternfly (Lycorma delicatula) establishment. Points are (R_0_, Z) pairings from a 120 x 120 rectangular grid across annual mean and amplitude temperature (h, g) parameter space (Fig. 3) for (a) diapause model variation (Pearson correlation coefficient: 0.74) and (b) non-diapause model variation (Pearson correlation coefficient: 0.82).

Different areas can exhibit similar *R*_0_ and *Z* values, but for different mechanistic reasons. In a large portion of the United States, including much of the Midwest, south, and mid-Atlantic regions, mean-temperature performance in diapause populations is tied primarily to development rates while noise inhibits performance by introducing diapause egg death in the winter (Fig. 8, Appendix 1: Point 3, noisy). In climates like that of much of California, the temperature trend is far above the egg cold mortality thresholds, and noise exerts a negative influence on performance through motile mortality in the spring and autumn (Fig. 8, 3b, Appendix 1: Point 7). In the arid climates of the U.S. southwest, such as that of Phoenix, AZ, summer heat mortality of motiles is the factor that both shapes mean temperature performance and—when exacerbated by noise—inhibits that performance. In Texas and the U.S. southeast, particularly parts of Florida, noise destabilizes populations by inducing mortality in motiles and decreasing egg-laying (Figs. 5b,c,e,f).

Climate change is expected to increase annual mean temperatures and decrease annual temperature amplitudes, meaning that the U.S. temperature area will shift downward and to the right in parameter space as climate change progresses (Di Cecco and Gouhier, 2018; Lyon et al., 2019; Olonscheck et al., 2021; Schneider et al., 2022). If the changes in the mean and amplitude are comparable, we infer that diapause populations in the upper region will experience an increase in resistance but little or no change in *R*_0_ (Fig. 4a, Fig 8). Some parts of southern California and the south may cross into the lower region and experience a fundamentally altered phenology with annual cycles of diapause and non-diapause populations, although *R*_0_ and *Z* may remain relatively unchanged.

Climatic variability has been shown in simple models to greatly raise or lower population growth rates depending on where within the environmental parameter space a population resides (e.g., Lawson et al., 2015). Specifically, change in population growth in the presence of environmental variability has typically been surmised through the convexity of the growth or vital rate function. Near average conditions that produce optimal performance, these functions often have a local maximum and are concave down, suggesting that variability lowers performance near optima. Conversely, functions have local minima and are concave up in environmental regimes in which performance is weakest, suggesting that variability should improve performance. Our model allows us to expand on this analysis by explicitly tracking instantaneous population change—determined by instantaneous vital rate responses to seasonal temperature trend and, importantly, noise—over the course of one year. The pattern of outcomes resulting from an ensemble of test realizations of noisy temperature profiles illuminates the effect of climatic variability on growth rates. The effects of variability are closely tied to the variation’s magnitude. If the standard deviation of the noise is sufficiently high, temperatures will be pushed to extremes and force high rates of mortality, regardless of the baseline temperature trend; this is why the resistance measure (Eq. (12)) is always well-defined.

At low noise standard deviations, the influence of noise in either increasing or decreasing growth rates is sensitive to the temperature trend, its relationship to the vital rate function parameters, and the nature of the vital processes themselves. Consider, for instance, the upper region in the diapause model (Fig. 3a) just below the diapause egg death threshold line (*h* – *g* = –3.96, Fig. 3b). Throughout this area, *R*_0_ varies up to its local optimum of ≈ 10, but *Z* remains very low (≤ 1°C) regardless of *R*_0_ (Fig. 4a). As the winter minimum temperatures in this area are extremely close to the diapause egg mortality threshold, any noise decreases growth factors, as expected. Consider again Detroit, MI (Fig. 8) where average temperature performance here is poor due to weak degree day accumulation. The same noise that might boost egg-laying rates at the end of the season also reliably negates those egg-laying gains by inducing winter egg mortality (Fig. 5a,c). While noise can stimulate both egg-laying and death, the latter often overwhelms the former. In contrast, further from the diapause egg death regime, along the SW boundary (Fig. 3a), the growth rate is still low, but low levels of noise are equally likely to drive growth rates greatly up or down. There are several U.S. urban areas with average temperature profiles that fall in this region and could see growth factors increase by over 50% of *R*_0_ with low levels of temperature variability (Fig. 9). Notably, several prominent wine-producing regions are represented, including locations across Napa and Sonoma Counties, CA, as well as the East End of Long Island, NY.

## 5. Summary and Conclusions

We developed and analyzed a stage-age-structured PDE model of the dynamics of spotted lanternfly (SLF) populations subjected to a stochastic temperature profile. The *R*_0_ patterns that emerge were dissected in terms of the relationship between the temperature parameters of climatic trend and those of SLF physiology. To understand growth under variable temperature conditions, we executed large ensemble simulations of the PDE with increasing stochastic noise *σ* and defined a new measure of resistance to temperature variability, *Z*, which is the largest *σ* for which the median annual growth factor remains above 1. We performed our simulations with two model variations, one in which SLF eggs can enter diapause to overwinter and another model variation in which SLF do not have the ability to diapause. Diapause tended to create rigid phenologies characterized by age-synchrony—univoltine in cooler climates but more complex in warmer ones—that can render a population either significantly more or less vulnerable to temperature fluctuations depending on the specific context. Non-diapause phenology is fundamentally different; there is a consistent presence of individuals across all life stages throughout the year—at least in warmer regions—the diversity of which protects the population against temperature fluctuations that may target specific subpopulations or developmental ages. Despite this, diapause populations have higher levels of overall resistance across the continental United States, even in the south, where non-diapause populations have high reproductive numbers. We also found that in certain mild climates where mean temperatures are well-separated from cold and heat mortality thresholds—as is the case in several U.S. winegrowing regions in California—realized growth at moderate, realistic levels of variability may be nearly twice the mean-temperature reproductive number.

Stakeholders should be aware that actual SLF growth can exceed predictions made by models that only look at average temperatures, but also models that include stochasticity without other processes like adaptation. While daily temporal patterns can cause outbreaks and declines that are uniform across regions, such climate-driven, macroscale variation of growth rate can interact with the adaptive potentials of populations and individual fitness (Belouard and Behm, 2023a). Furthermore, fitness varies at multiple scales due to landscape context (Ramirez et al., 2023). Large changes in annual growth make it likely that this pest can adapt quickly and require additional allocation of control resources to maintain target pest abundances within vineyards. Thus, understanding population resistance to temperature variability is complex, but allows for accurate forecasting that must be tested with broadscale monitoring data (De Bona et al., 2023) coupled to field-based individual tracking of pest fitness to estimate adaptation (Belouard and Behm, 2023b).

This manuscript is a first step in the exploration of the deeply nuanced sensitivity of SLF population dynamical behavior to stochastic temperature fluctuations, and the relationship between growth under average and variable temperature conditions. Here, the Langevin stochastic differential equation is used to formulate the noise component of the temperature model, in effigy of an AR(1) (autoregressive) model defined by parameters quantifying autocorrelation and noise magnitude. This formulation offers the advantage of simplicity and minimizes the size of the parameter space required to describe a noisy temperature profile in full. Knowing that noise magnitude likely varies spatiotemporally, this extension has the potential to provide an even more accurate description of establishment potential under variable conditions that agree with reality.

## Acknowledgments

We thank Jocelyn Behm, Nadège Bélouard, Stefani Cannon, Rujeko Chinomona, Alexis de la Cotte, Seba De Bona, Jason Gleditsch, Nick Huron, Kiera Kean, Anna Khan, Nour Khoudari, Shauna McManus, Sam Owens, Payton Phillips, Timothy Swartz, Jacob Woods, Mengsha Yao, and Nicole Zalewski for their comments on the work. Seba De Bona performed the interpolation and parameter fitting of temperature data. This work was funded by the United States Department of Agriculture (USDA) Animal and Plant Health Inspection Service Plant Protection and Quarantine under agreements AP20PPQS&T00C136, AP22PPQS&T00C146, AP22PPQS&T00C097, AP23PPQS&T00C090; the USDA National Institute of Food and Agriculture Specialty Crop Research Initiative Coordinated Agricultural Project Award 2019-51181-30014; the USDA Tactical Sciences for Agricultural Biosecurity Project Award 2022-68013-37139; and the Pennsylvania Department of Agriculture under agreements C9400000036, C94000833.

## Statement of Authorship

SML, BS, and MRH designed the study. SML developed the simulation software and conducted the analysis. BS and MRH provided guidance and supervision. SML and MRH wrote the manuscript. BS commented on the manuscript and provided edits

## Data and Code Availability

All data and code will be made available upon author request.

