## appendix 1 for "Quantifying population resistance to climatic variability: The invasive spotted lanternfly grape pest is buffered against temperature extremes in California"

List of video files (by order of appearance in the text):

- Point 1, no noise (point_1_nondiapause_no_noise.avi)
- Point 2 (point_2_nondiapause_no_noise.avi)
- Point 3, no noise (point_3_diapause_no_noise.avi)
- Point 4, no noise (point_4_diapause_no_noise.avi)
- Point 5 (point_5_nondiapause_no_noise.avi)
- Point 6 (point_6_diapause_noisy.avi)
- Point 1, diapause, noisy (point_1_diapause_noisy.avi)
- Point 1, non-diapause, noisy (point_1_nondiapause_noisy.avi)
- Point 4, noisy (point_4_diapause_noisy.avi)
- Point 3, noisy (point_3_diapause_noisy.avi)
- Point 7 (point_7_diapause_noisy.avi)

Point locations in parameter space:

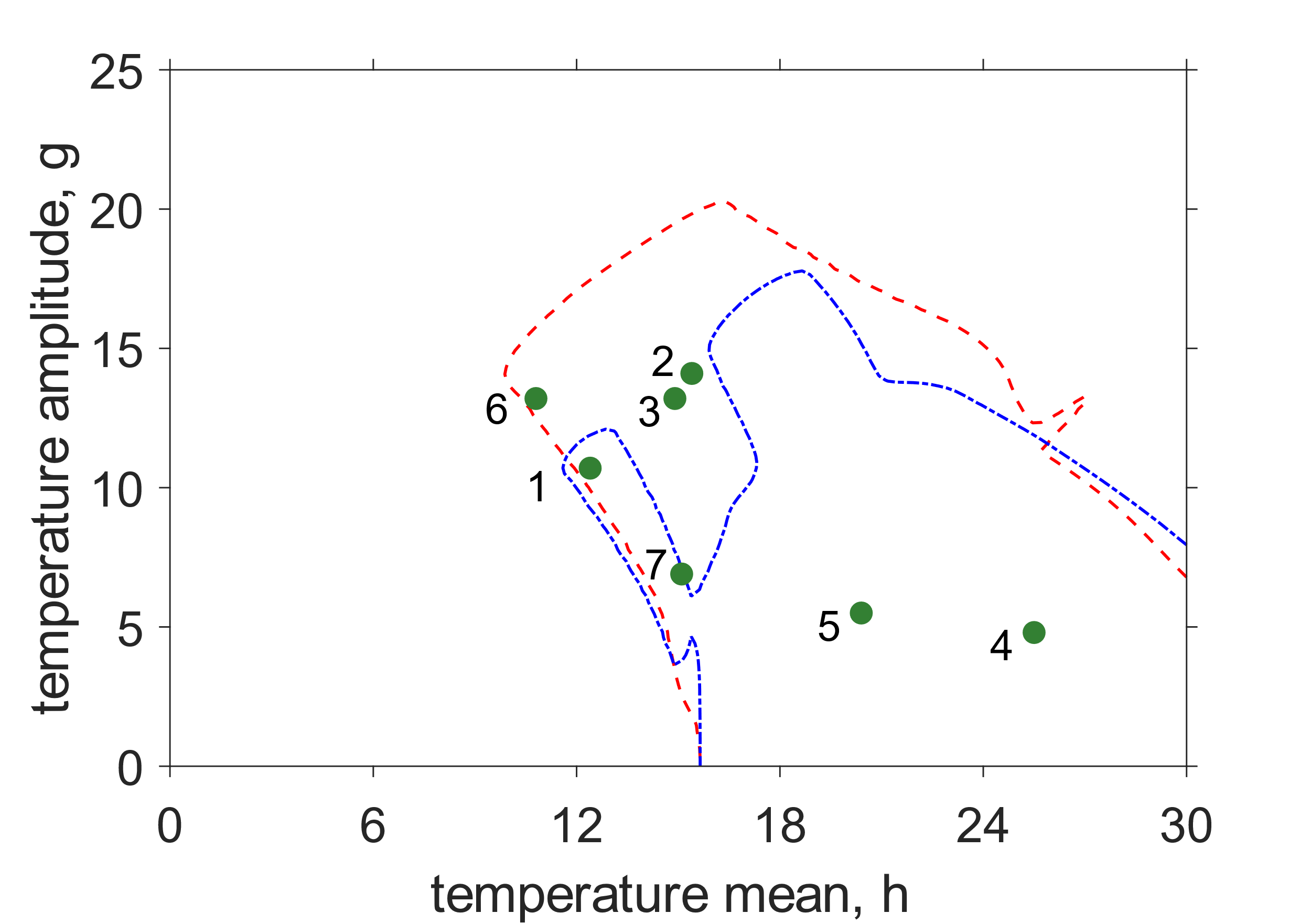

Details of each video example:

**Point 1**: Dynamics under deterministic temperature profiles in the “arm” region of the non-diapause model (see Fig. 3c)

- Filename: point_1_nondiapause_no_noise.avi
- Mean ($h$) = 12.4, Amplitude ($g$) = 10.7
- Diapause status: non-diapause model
- Description of simulation: A cohort of summer motiles develops and lays eggs in the fall. Most are still eggs when development ceases in early November, but about 20% have hatched into motiles by that point. Those that had already hatched die during the winter, while the eggs all survive and hatch in the spring.

**Point 2**: Dynamics under deterministic temperature profiles in the “gulf” region of the non-diapause model (see Fig. 3c)

- Filename: point_2_nondiapause_no_noise.avi
- Mean ($h$) = 15.4, Amplitude ($g$) = 14.1
- Diapause status: non-diapause model
- Description of simulation: At the start, there are cohorts of instars, egg-layers, and non-diapause developing eggs. The motiles pass through the egg-laying stage, laying a mass of eggs, all of which hatch well before development ceases in early November. All the newly-hatched motiles die during the winter, killing off the population entirely.

**Point 3, no noise**: Dynamics under deterministic temperature profiles in the upper region of the diapause model (see Fig. 3a)

- Filename: point_3_diapause_no_noise.avi
- Mean ($h$) = 14.9, Amplitude ($g$) = 13.2
- Diapause status: diapause model
- Description of simulation: At the start, an age-synchronized mass of motiles quickly lays a large cohort of eggs in diapause. The eggs overwinter in diapause, develop in the spring, and hatch in May.

**Point 4, no noise**: Dynamics under deterministic temperature profiles in the lower region of the diapause model (see Fig. 3a)

- Filename: point_4_diapause_no_noise.avi
- Mean ($h$) = 25.5, Amplitude ($g$) = 4.8
- Diapause status: diapause model
- Description of simulation: At the start, a small number of motiles—very spread out with respect to age—lays eggs in diapause throughout the fall. A few motiles remain after the winter solstice, and they lay non-diapause eggs, initiating rapid cycles of development and egg-laying through the non-diapause pathway. The large cohort of diapause eggs from the previous fall hatches in May, leading to a massive influx of motiles in the summer.

**Point 5**: Dynamics under deterministic temperature profiles in the lower region of the non-diapause model

- Filename: point_5_nondiapause_no_noise.avi
- Mean ($h$) = 20.4, Amplitude ($g$) = 5.5
- Diapause status: non-diapause model
- Description of simulation: Distributions of non-diapause developing and motiles—both spread out in age—persist throughout the year while the overall population size grows.

**Point 6**: Dynamics under noisy temperature profile near the intersection of southwest (“SW”, Fig. 3a) and northwest (“NW”) boundaries of upper region of the diapause model (see Fig. 7a,b)

- Filename: point_6_diapause_noisy.avi
- Mean ($h$) = 10.8, Amplitude ($g$) = 13.2, $\sigma$ = 2
- Diapause status: diapause model
- Description of simulation: An age-synchronized motile cohort develops and lays eggs in diapause in the fall, many of which die in the winter due to temperature decreases induced by noise. A small subset of the eggs survives and hatches the following spring, but the damage to the population is significant. (see Fig. 7b)

**Point 1, diapause, noisy**: Dynamics under noisy temperature profile near the middle of the southwest boundary (“SW”, Fig. 3a) of the upper region of the diapause model (see Fig. 7a,c)

- Filename: point_1_diapause_noisy.avi
- Mean ($h$) = 12.4, Amplitude ($g$) = 10.7, $\sigma$ = 2
- Diapause status: diapause model
- Description of simulation: A cohort of six motiles develops and lays approximately thirty-six eggs in the fall. There is no significant egg mortality in the winter, and after one year, 19 motiles remain (due to basal mortality). The one-year motile growth factor in this example surpasses three, which is much higher than the $R_{0}$ value at this point (~ 2), showing that noise significantly raised egg-laying rates. (see Fig. 7c)

**Point 1, non-diapause, noisy**: Dynamics under noisy temperature profile near the middle of the arm region (“arm”, Fig. 3c) of the upper region of the non-diapause model (see Fig. 7a,d)

- Filename: point_1_nondiapause_noisy.avi
- Mean ($h$) = 12.4, Amplitude ($g$) = 10.7, $\sigma$ = 3
- Diapause status: non-diapause model
- Description of simulation: A cohort of summer motiles develops and lays eggs in the fall, some of which hatch before development ceases. Those that hatch tend to die quickly, and those that remain in the egg stage mostly die off in the winter when noise brings temperatures below the non-diapause egg death threshold.

**Point 4, noisy**: Dynamics under noisy temperature profiles in the lower region of the diapause model (see Fig. 3a; compare with video for Point 4)

- Filename: point_4_diapause_noisy.avi
- Mean ($h$) = 25.5, Amplitude ($g$) = 4.8, $\sigma$ = 5
- Diapause status: diapause model
- Description of simulation: At the start, a small number of motiles—very spread out with respect to age—lays eggs in diapause throughout the fall. A few motiles remain after the winter solstice, and they lay non-diapause eggs, initiating rapid cycles of development and egg-laying through the non-diapause pathway. The large cohort of diapause eggs from the previous fall hatches in May, leading to a massive influx of motiles in the summer, many of which are promptly killed by heat induced by noise.

**Point 3, noisy**: Dynamics under noisy temperature profiles in the upper region of the diapause model (see Fig. 3a)

- Filename: point_3_diapause_noisy.avi
- Mean ($h$) = 14.9, Amplitude ($g$) = 13.2, $\sigma$ = 4
- Diapause status: diapause model
- Description of simulation: At the start, an age-synchronized mass of motiles quickly lays a large cohort of eggs in diapause. A large portion of the eggs are killed by low winter temperatures induced by noise, but about 25% survive, hatch, and develop.

**Point 7**: Dynamics under noisy temperature profiles in the lower part of the upper region of the diapause model.

- Filename: point_7_diapause_noisy.avi
- Mean ($h$) = 15.1, Amplitude ($g$) = 6.9, $\sigma$ = 6
- Diapause status: diapause model
- Description of simulation: A cohort of summer motiles develops and lays eggs, most of which survive the winter (a small amount—about 9%—are killed during a brief cold snap in December). The surviving eggs hatch in the spring and start to develop, but much of the surviving population is killed off by cold snaps induced by noise that bring temperatures below the motile cold mortality threshold.
